# Pan-cancer analysis reveals context-dependent roles of LINE-1 ORF1p in immune regulation and copy number alterations

**DOI:** 10.1101/2025.10.03.680361

**Authors:** Efiyenia Ismini Kaparos, Wenjing Zhang, Ryusei Miyanaga, Wilson McKerrow, Jef Boeke, Teresa Davoli, Paolo Mita, Kelly V. Ruggles, David Fenyö

**Affiliations:** Institute for Systems Genetics, New York University Grossman School of Medicine, New York, NY, USA; Department of Biochemistry and Molecular Pharmacology, New York University Grossman School of Medicine, New York, NY, USA; Division of Precision Medicine, Department of Medicine, New York University Grossman School of Medicine, New York, NY USA

## Abstract

L1 comprises 17% of the human genome, with 50-150 full-length sequences capable of retrotransposition. Although largely inactive in normal somatic tissues, tumorigenesis leads to L1 derepression and overexpression of its RNA binding chaperone protein, ORF1p. A potential cancer biomarker, ORF1p expression is a hallmark of multiple cancers and an early event in precursor lesions. Our study provides a comprehensive pan-cancer analysis of ORF1p using CPTAC proteogenomic data, revealing a dichotomous role in modulating immune responses. Integrated analyses and supervised learning models divide ORF1p-high tumors into two groups: (1) high ORF1p associates with immunosuppression, reduced interferon signaling and diminished immune cell infiltration (HNSCC and LSCC), and (2) ORF1p-high tumors associate with immune activation (UCEC). Linear regression models reveal that cancer-specific aneuploidies may underlie this immune dichotomy, highlighting the prognostic significance of ORF1p and informing new strategies to leverage detection of plasma-circulating ORF1p to enhance immunotherapeutic efficacy in different tumor contexts.

## INTRODUCTION

Transposable elements comprise about half of the human genome, with retrotransposons accounting for just over 40%.^1,2^ Among them, the long interspersed nuclear element-1 (LINE-1 or L1) makes up 17% of these sequences and is the only active autonomous element.^1^ Although most of the ∼500,000 L1 copies in the human genome are truncated or mutated, about 50–150 copies remain full-length with intact open reading frames (ORF).^3–5^ These elements retrotranspose through a “copy-and-paste” mechanism, in which L1 RNA is reverse transcribed and integrated into new genomic sites.^6,7^ This process requires the two L1 encoded proteins: ORF1p, an RNA-binding protein, and ORF2p, which provides endonuclease and reverse transcriptase functions and also capable to drive the mobilization of non-autonomous repeats such as SINEs.^8–11^ Together with L1 RNA, ORF1p and ORF2p form ribonucleoprotein particles (RNPs) that mediate nuclear entry and integration.

Due to its potentially mutagenic nature, L1 is suppressed in most normal somatic tissues.^12,13^ Host cells have evolved multiple defense mechanisms to regulate and restrict L1 activity, but in many forms of cancer L1 repression mechanisms are lost and repeat elements are often derepressed.^14–20^ Pattern recognition receptors (PRR) recognize derepressed L1 byproducts triggering type I interferon (IFN) responses similar to the one activated by exogenous retrovirus invasions. ^21–23^

Holistically understanding the functional role of L1 derepression in cancer requires an examination of L1 activity, defined here as its ability to re-insert into new random genomic loci, as well as the consequences of L1 expression across its lifecycle, including its RNA regulation, ORF1p/ORF2p expression and L1 RNP interaction with the tumorigenic cellular context.^24^ Current *in vivo* and *in vitro* studies investigating L1 in cancer have relied on engineered L1 overexpression systems and reporter assays that miss physiologically relevant contexts such as the complete tumor ecosystem, comprised of cancer-associated fibroblasts and immune system interactions.^25–31^ Despite the extensive genomic characterization of L1 activity in cancer^32–35^, comprehensive pan-cancer proteomic analyses of L1 ORF1p aimed at understanding the consequences of its overexpression in neoplasia are still limited.^36,37^ Our study aims to fill this gap by characterizing ORF1 protein expression, both because its expression is more easily detectable by proteomic approaches compared to ORF2p^38^ and because recent studies have highlighted a complex role for ORF1p in tumorigenesis.^25,39–42^ In addition to the establishment of ORF1p as a potential diagnostic marker for multiple cancers, a study in pancreatic cancer identified an unexpected feed-back loop in which ORF1p, through its mRNA binding activity, may decrease the IFN-I response triggered by its own L1 mRNA.^42^ To better understand the impact of L1 ORF1p derepression in *in vivo* tumor contexts, we used the NCI-supported Clinical Proteomic Tumor Analysis Consortium’s (CPTAC) harmonized pan-cancer dataset (n = 1,043) spanning 10 cancer types^43^, and integrated L1 quantifications with proteogenomic data with emphasis on L1 ORF1p. In our analysis, we grouped patients based on their ORF1p protein expression levels and applied supervised learning to map downstream effects on gene, protein, and phosphoprotein expression, as well as immune-related changes and copy number alterations. We identified tissue specific consequences of L1 expression in the cellular adaptation to high ORF1p expression and in the immune response triggered by L1 derepression and describe molecular vulnerabilities unique to persistent ORF1p overexpression with prognostic potential.

## RESULTS

### Pan-cancer characterization of L1 ORF1p

We quantified L1 RNA and ORF1p in all 10 CPTAC tumor types and identified novel L1 somatic insertions where whole genome sequencing (WGS) was available (Figures 1A and S1A) as previously described.^44^ Of the 1,043 samples with transcriptomic, proteomic or WGS data available, we identified 929 tumors with at least one L1 measurement detected (Figure S1B). This included BRCA, CCRCC, COAD, OV, and UCEC tumors.^44,45^ Out of the 929 tumors in our analysis, 808 included ORF1 protein quantification (Figures 1B and S1C), with the highest median ORF1p expression identified in UCEC and the lowest in BRCA (Figure S1D). 776 samples in our dataset had L1 RNA quantification (Figures 1B and S1C), with the highest median L1 RNA expression in HNSCC and the lowest in GBM (Figure S1D). In GBM tumors, we detected ORF1p abundance in only 8% of samples (Figure S1E), consistent with previous findings showing that L1 retrotransposition and ORF1p expression are infrequent in aggressive brain tumors.^40,46,47^ Thus, we have excluded GBM from all proteomic-based analyses in this study.

**Figure 1.**
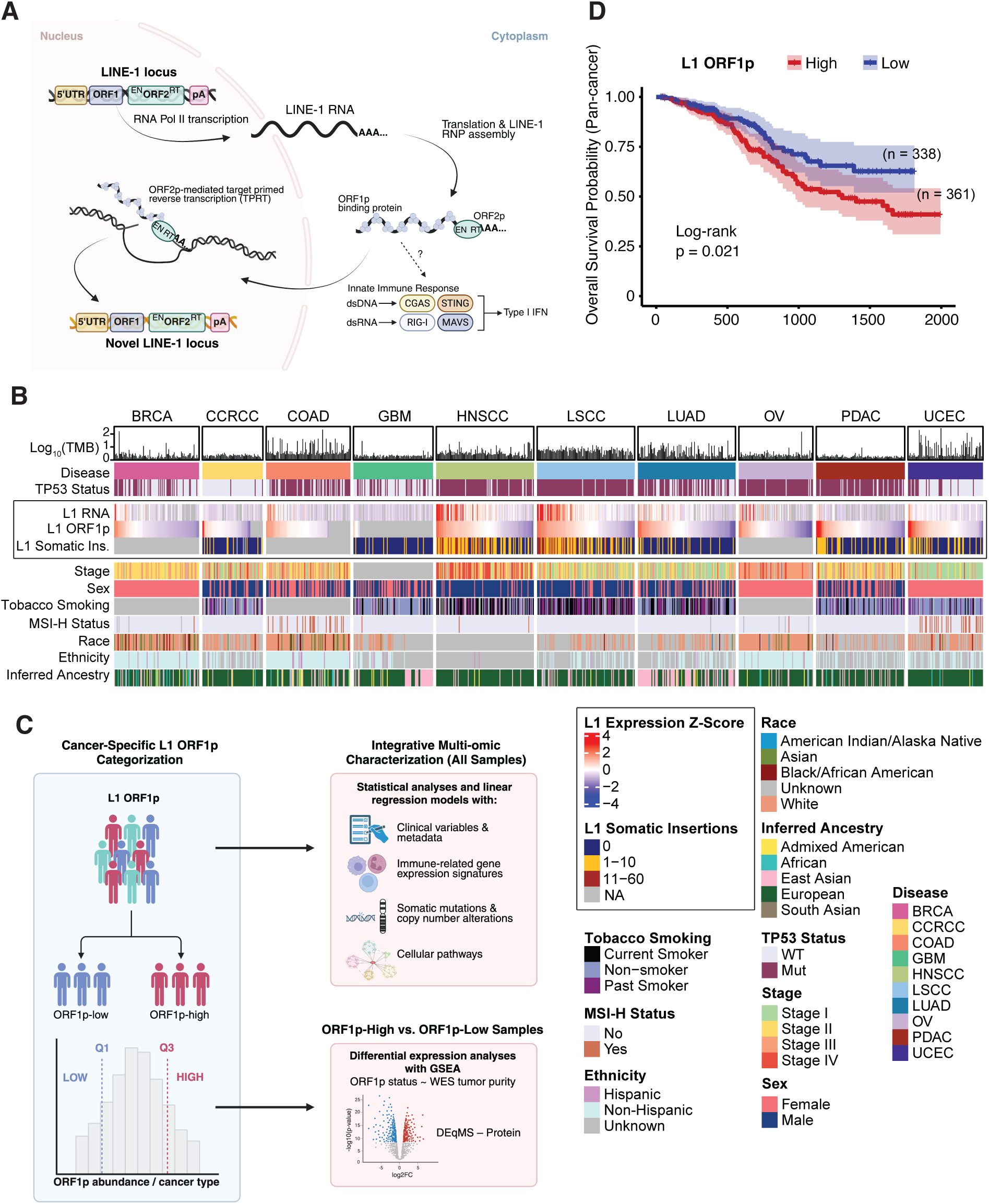
Pan-cancer overview of L1 activity. (A) Schematic of L1 life cycle. L1 consists of two open reading frames, ORF1 and ORF2, and is transcribed by the cell’s machinery. L1 mRNA translocates to the cytoplasm for ORF1/2 translation and ribonucleoprotein assembly whereby it re-enters the nucleus for ORF2p-mediated target primed reverse transcription (TPRT) into a new genomic locus. Formation of L1 dsRNA from L1 mRNA and DNA:RNA hybrids and L1 cDNA from reverse transcription can be sensed by cytosolic DNA and RNA sensors to trigger an innate immune response and type I IFN induction. (B) Heatmap ordered by increasing ORF1p expression stratified by CPTAC cancer type profiling clinical variables. L1 quantifications are indicated by the black box on the heatmap and legend. (C) Schematic outlining ORF1p-high and ORF1p-low sample categorization and downstream proteogenomic analysis. (D) Pan-cancer Kaplan-Meier plot comparing overall survival (OS) for patients stratified by ORF1p status, log rank test. Numbers in parentheses represent the pan-cancer sample size for each group.

Except for CCRCC, L1 RNA and ORF1p correlated strongly (Spearman *R* > 0.4) and significantly (*P* value < 0.001) across cancer types, suggesting that L1 ORF1p level is mainly dependent on transcription and less on post-transcriptional regulations (Figures 1B and S1F). Incidence of novel somatic L1 insertions correlated most strongly with L1 RNA expression (Spearman *R* > 0.6, *P* value < 10^-12^) in HNSCC and LSCC tumors (Figure S1G), which also had the greatest volume of insertions ranging from 11-60 (Figure 1B). These findings suggest that HNSCC and LSCC are highly L1 active tumors, as previously shown^32,33^, and may indicate an enhanced capacity of these tumors to sustain elevated L1 expression and activity. This observation could be at least partially explained by the fact that the vast majority of HNSCC and LSCC tumors in these cohorts harbor p53 mutations (Figures 1B and S1H), a known pre-requisite for L1 expression.^20,44^ On the other hand, our measurements also show similar p53 mutation frequencies in PDAC and OV compared to HNSCC and LSCC tumors, suggesting that additional regulatory mechanisms beyond p53 influence L1 expression. Overall, these results confirm our ability to quantify L1 at various points of its lifecycle (RNA, ORF1 protein and novel insertions) (Figure 1A) and demonstrate that L1 is derepressed across all CPTAC tumor types (Figures 1B).

To characterize L1 ORF1p expression in cancer, we first grouped samples as ORF1p-high (top quartile) or ORF1p-low (bottom quartile) and assessed the association of this grouping with multimodal molecular features, cancer types, subtypes, and clinical outcomes (Figure 1C). Past studies have shown an association between L1 RNA expression and methylation status and poor outcomes across several cancer types.^48,49^ To assess the relationship between L1 protein expression and survival in CPTAC, we stratified our samples by ORF1p and L1 RNA expression, grouping samples into L1 high and L1 low (Methods and Figure 1C). Using pan-cancer Kaplan–Meier analysis, we found significantly lower survival for ORF1p-high patients compared to ORF1p-low across CPTAC cohorts (log-rank *P* = 0.021, Figures 1D and S1I). Parallel L1 RNA analysis revealed no significant difference in overall survival between L1 RNA-high and L1 RNA-low patients across cancer types, suggesting that L1 ORF1p overexpression has a functional role that affects cancer outcomes independent of L1 RNA expression (Figure S1J). Survival analyses within individual cancer types did not yield any significant differences in overall survival based on ORF1p expression, possibly due to small sample sizes (Figure S1K).

### Clinical associations with ORF1p expression

We evaluated the association of ORF1p expression with clinical and demographic variables (Figure 2A). Interestingly, L1 ORF1p expression was significantly increased in current smokers (P = 0.006) and past smokers (P = 0.015) when compared to non-smokers in HNSCC, with no significant differences based on smoking identified in other cancer types. LUAD patients with East Asian inferred ancestry had significantly higher (P value = 0.047) ORF1p levels compared to those with inferred European and African ancestry.

**Figure 2.**
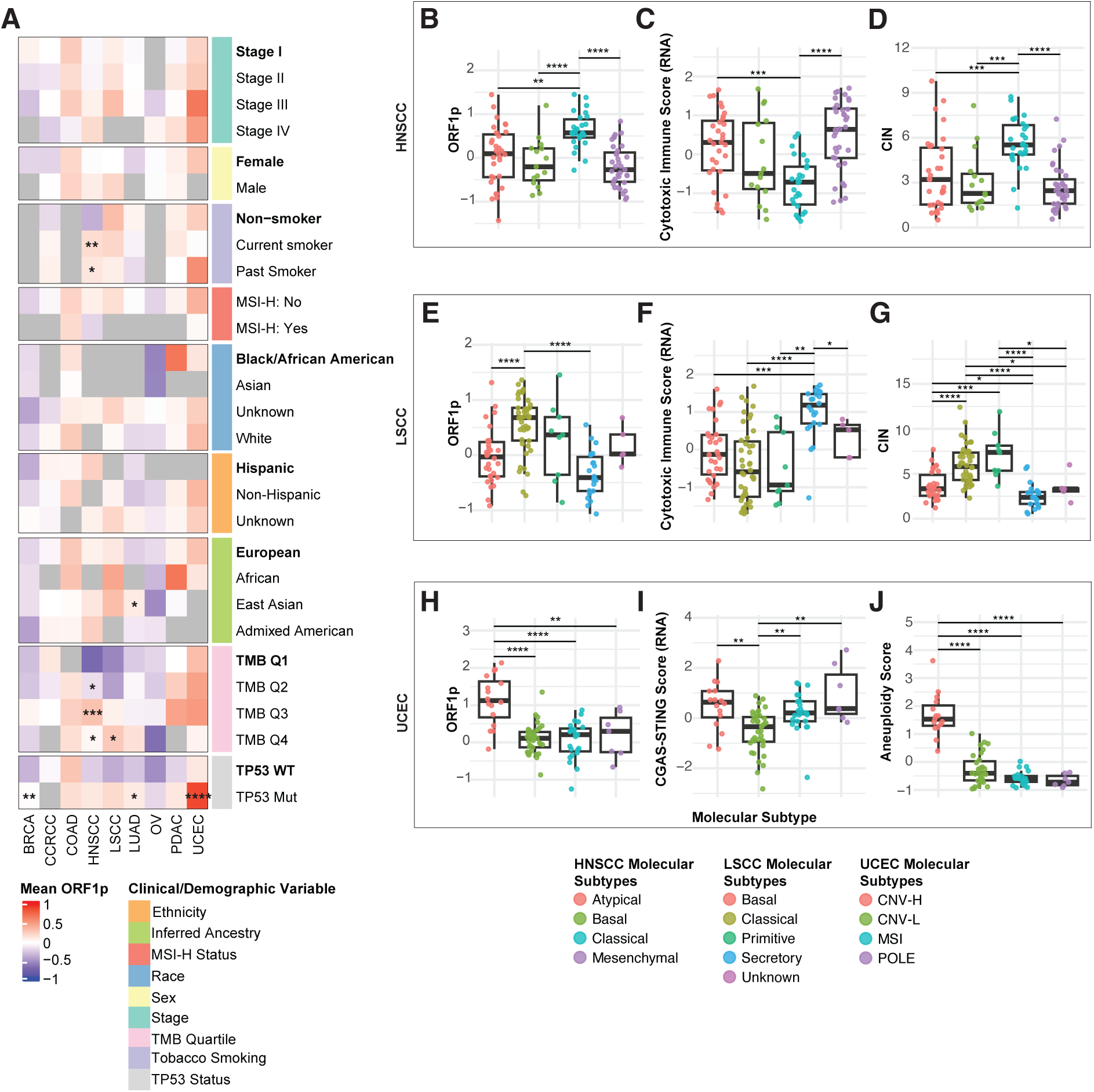
Clinical variable and tumor molecular subtype associations with L1 ORF1p. (A) Mean ORF1p expression stratified by clinical variable sub-categories. Rows are sub-categories grouped by clinical variables. Columns are cancer types. Bolded sub-categories indicate the reference group used for Wilcoxon rank sum test of ORF1p distribution with BH p-value correction for variables with more than 2 sub-categories for each clinical variable. Significant comparisons indicated with asterisk (*). (B-G) Boxplots of (B,E) ORF1p, (C,F) cytotoxic immune score and (D,G) CIN stratified by molecular subtypes of HNSCC and LSCC, respectively. (H-J) Boxplots of (H) ORF1p, (I) CGAS-STING score and (J) aneuploidy score stratified by molecular subtypes of UCEC. P ≤ 0.05, **p≤ 0.01, ***p ≤ 0.001, ****p ≤ 0.0001.

Tumor mutational burden (TMB), defined as the number of somatic mutations per megabase of a genomic sequence, is believed to generate immunogenic neoantigens that enhance patient response to immune checkpoint inhibitors (ICI).^50^ To determine associations of ORF1p with TMB in our cohorts, we categorized TMB into groups using quartiles from the TMB distribution per cohort (Q1, Q2, Q3 and Q4). ORF1p expression was found to be significantly higher in tumors with higher TMB levels in HNSCC (P = 0.026 for TMB Q2, 0.00013 for TMB Q3 and 0.027 for TMB Q4) and LSCC (P = 0.012 for TMB Q4). HNSCC and LSCC are tumor types characterized by high TMB^50^, suggesting that L1 ORF1p expression is positively associated with an increasing TMB in squamous cell carcinomas (Figure 2A). Though L1 RNA and ORF1p expression are significantly higher in p53 mutant tumors compared to p53 WT tumors at the pan-cancer level (Figure S2A-S2B), significant differences between ORF1p expression and p53 mutation status at the cohort-specific level were detected only in BRCA, LUAD and UCEC (Figure 2A), while LSCC was the only cohort to have a significant difference in L1 RNA expression and p53 mutation status (Figure S2C). Though lack of significance at the cohort level is largely due to sample size, there may be additional p53 independent mechanisms of regulation for the other tumor types. No significant differences in L1 ORF1p expression were found for stage, sex, MSI-high (MSI-H) status, race, and ethnicity (Figure 2A). Parallel analyses with L1 RNA resulted in similar significant trends seen in the ORF1p analysis, with higher L1 RNA expression correlating with increased TMB levels in HNSCC and LSCC (Figure S2C). Current smokers of HNSCC had significantly higher L1 RNA levels than non-smokers (*P* = 0.0074), while current smokers of UCEC had significantly lower L1 RNA expression than non-smokers (*P* = 0.036), a trend also seen at the L1 protein level as well (Figures 2A and S2C).

### ORF1p expression in tumor subtypes

We then assessed ORF1p expression by CPTAC genomic molecular subtypes to determine whether ORF1p expression can be utilized as an additional patient stratification metric (Figures 2B-J and S2D-L).^51–59^ HNSCC, LSCC and UCEC showed significant differences in ORF1p expression across molecular subtypes, with the highest ORF1p expression observed in the classical subtypes in HNSCC and LSCC, respectively, and the CNV-H subtype in UCEC compared to all other subtypes. The lowest expression of ORF1p was found in mesenchymal and secretory subtypes of HNSCC and LSCC, respectively (Figures 2B, 2E, and 2H, P < 0.001). Similarly, the chromosomal instability (CIN) genomic subtype of COAD had significantly higher ORF1p expression than the mesenchymal subtype (Figure S2F).

High ORF1p-expressing classical subtype tumors in HNSCC and LSCC are known to have similar molecular profiles, including *TP53* mutation, strong smoking history, low immune activation, and high CIN.^55,58,60^ We found that tumors in these subtypes also had significantly lower cytotoxic immune scores compared to other molecular subtypes (Figures 2C and 2F) and significantly higher CIN (Figures 2D and 2G). Low ORF1p- expressing mesenchymal and secretory subtypes of HNSCC and LSCC were characterized by upregulation of immune-related pathways based on cytotoxic immune score (Figures 2C and 2F). Similar to HNSCC and LSCC, ORF1p high-expressing CIN- subtype tumors of COAD had significantly lower cytotoxic immune score levels compared to the mesenchymal subtype (*P* < 0.01), along with higher CIN (*P* < 0.01) as expected and significantly higher aneuploidy scores compared to the MSI subtype (*P* < 0.0001) (Figure S2F).

UCEC CNV-H tumors are characterized by frequent copy number alterations and increased expression of cell cycle and antiviral response proteins.^54,61^ We found that these tumors had the highest levels of CIN compared to all other UCEC subtypes, similar to the classical subtype of HNSCC and LSCC (Figure 2J; *P* < 0.001). Although there were no significant differences between UCEC subtypes for cytotoxic immune score (Figure S2L), CNV-H tumors had significantly higher CGAS-STING scores, representing antiviral signaling activation, than CNV-L tumors (Figure 2I) and the highest aneuploidy scores when compared to all other molecular subtypes of UCEC, as expected (Figure S2L).

Collectively, these data demonstrate that ORF1p expression can distinguish between more and less aggressive molecular subtypes within each cancer type, with high ORF1p expression consistently associated with high CIN and reduced immune response in HSCC, LSCC and COAD and, in contrast, associated with increased immune response in UCEC.

### ORF1p modulates the expression and post-translational modification of proteins involved in IFN response

Previous work in pancreatic cancer showed that repetitive elements, including L1, are able to trigger IFN-I activation through “viral mimicry” processes.^25,29,42^ *In vitro* and *in vivo* studies identified cytoplasmic dsRNA, RNA:DNA hybrids or dsDNA as potential molecules produced by retrotransposons expression and activity and able to trigger and IFN-I response.^23,62–65^ Moreover, the L1 ORF1p protein itself was shown to “mask” dsRNA L1 adducts decreasing the activation of IFN-I mediated responses, inducing an opposite effect from the one triggered by L1 mRNA expression.^42^

To gain a pan-cancer view of the L1 involvement in viral mimicry pathways during tumorigenesis and gain insights on the molecular details at play in *in vivo* contexts, we first conducted differential proteomic expression analysis accounting for tumor purity between ORF1p-high and ORF1p-low samples for each cancer type (Methods), identifying tumor specific and pan-cancer associations (Table S1, Figure S3A-I). After filtering for proteins with an adjusted *P* < 0.2 in at least four of the cohorts, we identified a common signature of 17 proteins differentially expressed in ORF1p-high samples (Figure 3A). Of the 17 proteins identified 10 are directly connected to RNA metabolism and/or ribonucleoprotein metabolism (Figure 3A, bold), suggesting that RNA-linked processes may be key for the tissue and tumor response to ORF1p expression. This is perhaps not surprising considering the well-known function of ORF1p as RNA binding protein.^66,67^ Specifically, the ORF1p proteomic signature includes reduced expression of selenium binding protein 1 (SELENBP1) and vacuolar protein sorting VPS13C and increased expression of transcription factor TFB2M, nuclear export receptor XPO5, replication factor RFC4 and protein phosphatase Mg2+/Mn2+ dependent 1G (PPM1G) (Figure 3A). Several of these proteins are established repressors of IFN signaling, inflammation or viral defense processes^68–70^, suggesting that tumors may actively attempt to limit the L1-triggered IFN response (Table 1).

**Figure 3.**
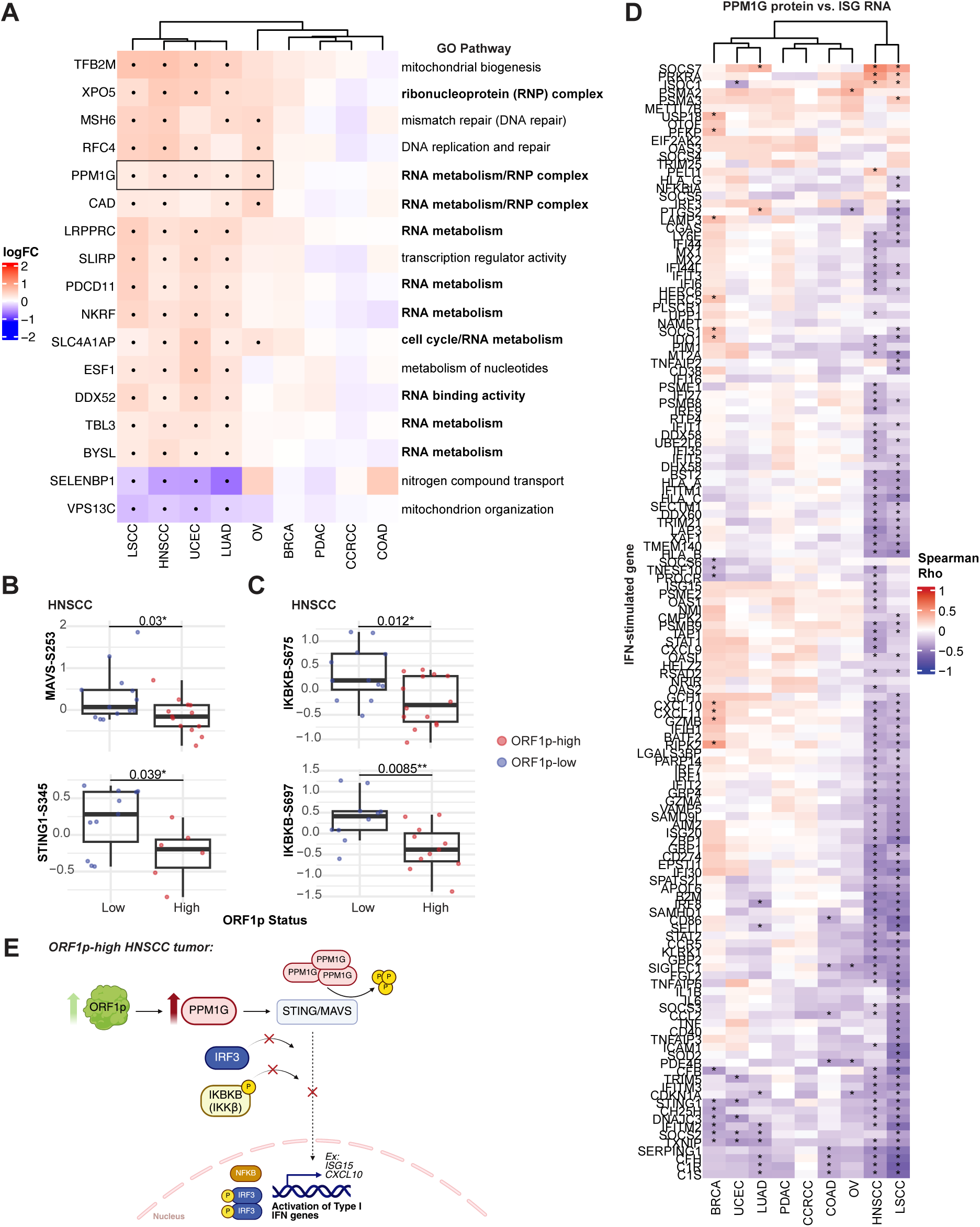
ORF1p modulates the expression and post-translational modification of proteins involved in IFN response. (A) Heatmap of logFC for differentially expressed proteins in ORF1p-high tumors filtered for P adjusted < 0.2 in at least four cohorts indicated by black dot (•). Associated Gene Ontology (GO) pathway to the right of heatmap with RNA/RNP metabolism pathways bolded. PPM1G is boxed to denote further investigation in panels B-D. (B-C) Protein-normalized phosphosite abundance of immune genes downstream of PPM1G significant in ORF1p-high and ORF1p-low groups of HNSCC. Two-tailed T-test. (D) Spearman correlation of RNA abundance of type I IFN genes from IFN signature gene set (Methods) with PPM1G protein in HNSCC. (E) Proposed model summarizing findings from panels A-D explaining post-translational control of immune suppression in ORF1p-high tumors of HNSCC. For T-tests: p ≤ 0.05, **p≤ 0.01, ***p ≤ 0.001, ****p ≤ 0.0001.

**Table 1.** ORF1p-high proteomic signature from differential protein expression.

| Protein | Known function | Direct involvement in IFN response/inna te immunity: | Indirect involvement in IFN response/inna te immunity: | Activator or repressor of IFN response or innate immunity? | Notes | References (PMID) |
| --- | --- | --- | --- | --- | --- | --- |
| TFB2M | Mitochondrial transcription factor B2; essential for mitochondrial DNA transcription initiation | | YES | Activator? | In ovarian cancer, TFB2M overexpression increases cytosolic mtDNA. This activates the TLR9/NF-κB/IL-6 pathway. | PMID: 35577351 |
| XPO5 | Exportin-5; mediates nuclear export of pre-miRNAs and some proteins |  | YES | Repressor | XPO5 can regulate interferon response by regulating the microRNA (miRNA) pathway, which is a key component of the innate immune system. Some viruses, such as SARS-CoV-2, can target and degrade XPO5 to inhibit the maturation of miRNAs. XPO5 also binds dsRNA and stimulates RBP and ADAR1 export from the nucleus. | PMID: 19124606<br>PMID: 14631048 |
| MSH6 | DNA mismatch repair protein; also functions in antibody somatic hypermutation and class switch recombination |  | YES | Repressor? | Reduced expression of the DNA mismatch repair (MMR) protein MSH6 is associated with a more active immune response and a stronger interferon signature in several cancers, particularly endometrial cancer. This inverse relationship suggests that MSH6's normal function may suppress immune signaling, and its loss can trigger an immune response that is more sensitive to immunotherapy. | PMID: 38390329 |
| RFC4 | Replication factor C subunit; involved in DNA replication and repair | NO | NO |  |  |  |
| PPM1G | Protein phosphatase; regulates pre-mRNA splicing and chromatin structure | YES |  | Repressor | Known regulator of the innate immune adaptor proteins stimulator of interferon response cGAMP interactor 1 (STING1) and mitochondrial antiviral signaling protein (MAVS). | PMID: 33219031 |
| CAD | Multifunctional enzyme in de novo pyrimidine biosynthesis (carbamoyl-phosphate synthase 2, aspartate transcarbamylase, dihydroorotase) | | YES | Repressor | Carbamoyl-phosphate synthetase 2, aspartate transcarbamylase, and dihydroorotase (CAD) influences the interferon (IFN) response through its role in pyrimidine synthesis and through complexing with innate immune proteins. CAD also deamidates the NF-κB pathway transcriptional factor RelA (also known as p65). | PMID: 38743960 |
| LRPPRC | Leucine-rich pentatricopeptide repeat-containing protein; regulates mitochondrial gene expression and mRNA stability | YES |  | Repressor | LRPPRC interacts with and inhibiting the mitochondrial antiviral signaling protein MAVS, a key adaptor protein in antiviral immunity. | PMID: 30070380 |
| SLIRP | Stem-loop interacting RNA binding protein; mitochondrial RNA stability and regulation | YES |  | Activator | SLIRP amplifies the interferon (IFN) response by stabilizing and promoting the release of mitochondrial double-stranded RNAs (mt-dsRNAs), which then activate MDA5, a pattern recognition receptor. | PMID: 40253699 |
| PDCD11 | Programmed cell death protein 11; nucleolar protein involved in rRNA processing and NF-κB pathway regulation | | YES | Repressor | PDCD11 does not have a well-documented role in directly regulating the interferon (IFN) response. However, there is evidence of an indirect, regulatory connection through its effect on nuclear factor-kappa B (NF-κB). PDCD11 interacts with ADAR1. | PMID: 39673305<br>PMID: 32709934 |
| NKRF | NF-κB repressing factor; transcriptional repressor of NF-κB target genes including IFN-β and cytokines | YES | | Repressor | NKRF plays a key role in the interferon (IFN) response by inhibiting the transcription of certain genes, including the IFN-β gene. NKRFs function is to prevent an excessive or prolonged immune reaction that could harm the host. | PMID: 37084822 |
| SLC4A1AP | Solute carrier family 4A1 adaptor protein; involved in centrosome duplication and mitotic spindle formation | NO | NO |  |  |  |
| ESF1 | ESF1 homolog; nucleolar protein required for pre-rRNA processing | NO | NO |  |  |  |
| DDX52 | DEAD-box helicase 52; functions in ribosome biogenesis and RNA processing |  |  | Repressor? | DDX52 appears to act as an antiviral protein against some viruses, but there is no direct evidence that DDX52 is part of a canonical interferon-response pathway. | PMID: 29146961 |
| TBL3 | Transducin beta-like 3; WD40 repeat protein involved in RNA metabolism | YES |  | Repressor | TBL3 is an interferon-stimulated gene (ISG) whose expression increases during an interferon antiviral response and that inhibits IRF1 (interferon (IFN) regulatory factor-1). | PMID: 25051498 |
| BYSL | Bystin-like protein; ribosome biogenesis and early embryonic cell adhesion | NO | NO |  |  |  |
| SELENBP1 | Selenium-binding protein 1; implicated in redox regulation, oxidative stress response, and cancer biology |  | YES? | Repressor?/Activator? | SELENBP1 is involved in regulating the inflammatory and oxidative stress pathways that are modulated by the interferon response. Its exact role is complex and may vary depending on the context, such as in autoimmune diseases versus cancer. | PMID: 36253721<br>PMID: 38172565 |
| VPS13C | Vacuolar protein sorting 13 homolog C; lipid transfer protein at organelle contact sites, mutations linked to Parkinson's disease | YES |  | Repressor | Loss of the lipid-transport protein VPS13C activates the innate immune interferon response through the cGAS-STING pathway. This occurs because dysfunctional lysosomes, caused by a lack of VPS13C, become leaky, leading to the escape of mitochondrial DNA (mtDNA) into the cytosol. | PMID: 35657605 |

In particular PPM1G is a known regulator of the innate immune adapter protein STING1 (STimulator of INterferon response cGAMP interactor 1) and MAVS (Mitochondrial AntiViral Signaling) protein.^69^ To determine whether increased expression of PPM1G in ORF1p-high tumors also results in changes of phosphorylation of its downstream targets STING and MAVS, we leveraged CPTAC’s phosphoproteomic data and assessed differences between ORF1p-high and ORF1p-low groups for all STING and MAVS phosphosites (Figure 3B and Table S2). Due to overexpression of PPM1G in most cancer types, we expected to find decreased phosphorylation of STING and MAVS in ORF1p-high tumors. Indeed, ORF1p-low samples in HNSCC had significantly higher phosphorylation of MAVS-S253 (*P* = 0.03) and STING1-S345 (*P* = 0.039) compared to ORF1p-high samples. No other cancer types exhibited significant differences in phosphorylation of PPM1G targets in this direction (Table S2). To determine if dephosphorylation of STING and MAVS affects downstream activation of the IFN program^71,72^, we evaluated the phosphorylation status of inhibitor of nuclear factor kappa B kinase subunit beta (IKBKB) across cancer types and found reduced phosphorylation of IKBKB in HNSCC ORF1p-high samples at phosphosites Ser675 and Ser697 (Figure 3C and Table S2). Phosphorylation of IKBKB triggers phosphorylation of the inhibitor that silences the NF-𝜿B complex^73,74^, and consistent with this relationship, expression of downstream IFN-stimulated genes was found to be significantly negatively correlated with PPM1G protein expression in HNSCC and LSCC (adjusted *P* value < 0.05). Tumor types with the highest levels of ORF1p, HNSCC and LSCC, clustered together in their IFN expression signature (Figure 3D). OV and COAD tumors also harbored few significant negative correlations between IFN-stimulated gene expression and PPM1G protein expression (Figure 3D).

Together, these findings indicate that ORF1p-high HNSCC and LSCC tumors upregulate proteins to reduce and control IFN-I signaling. One such protein is PPM1G, a phosphatase whose upregulation correlates with dephosphorylation of MAVS, STING and IKBKB proteins, negatively modulating the IFN-I and NF-𝜿B innate immune response (Figure 3E). This mechanism was significantly observed in HNSCC tumors (Figures 3B-C), likely facilitating tolerance to L1 overexpression.

### ORF1p expression regulates immune pathways and immune cell infiltration in a tumor-specific manner

Despite the established role of IFN mediated inflammation in cancer development, studies focused on the consequences of the L1-mediated IFN activation on immune cell recruitment and activity are still limited.^75,76^ To more systematically investigate the role of ORF1 protein expression in immune regulation, we performed gene set enrichment analysis (GSEA) on differentially expressed proteins in ORF1p-high tumors to assess ORF1p-associated pathway dysregulation for each cancer type (Table S3). Previous studies have reported enrichment in pathways of DNA repair and cell cycle regulation in ORF1p-high tumors.^44^ Consistent with this, we found that in all tumor types except for CCRCC, pathways of oxidative phosphorylation, glycolysis, and MYC and MTORC1 signaling were significantly enriched in tumors with high ORF1p-expressing tumors (Figure 4A). Further, we found that pathways involving innate immune and inflammatory signaling, hypoxia, epithelial mesenchymal transition (EMT), apoptosis and myogenesis were downregulated in ORF1p-high LSCC, PDAC, COAD and HNSCC tumors (Figure 4A). These findings align with previous studies that have demonstrated a negative association of immune response with L1 retrotransposition.^29,35,42^ Interestingly and more specifically, we also noticed that several IFN Hallmark pathways were upregulated in ORF1p-high for CCRCC, OV, UCEC and BRCA that, in this context, seem to behave in an opposite manner compared to LSCC, PDAC, COAD and HNSCC as for our previous results (Figure 4A, bold).

**Figure 4.**
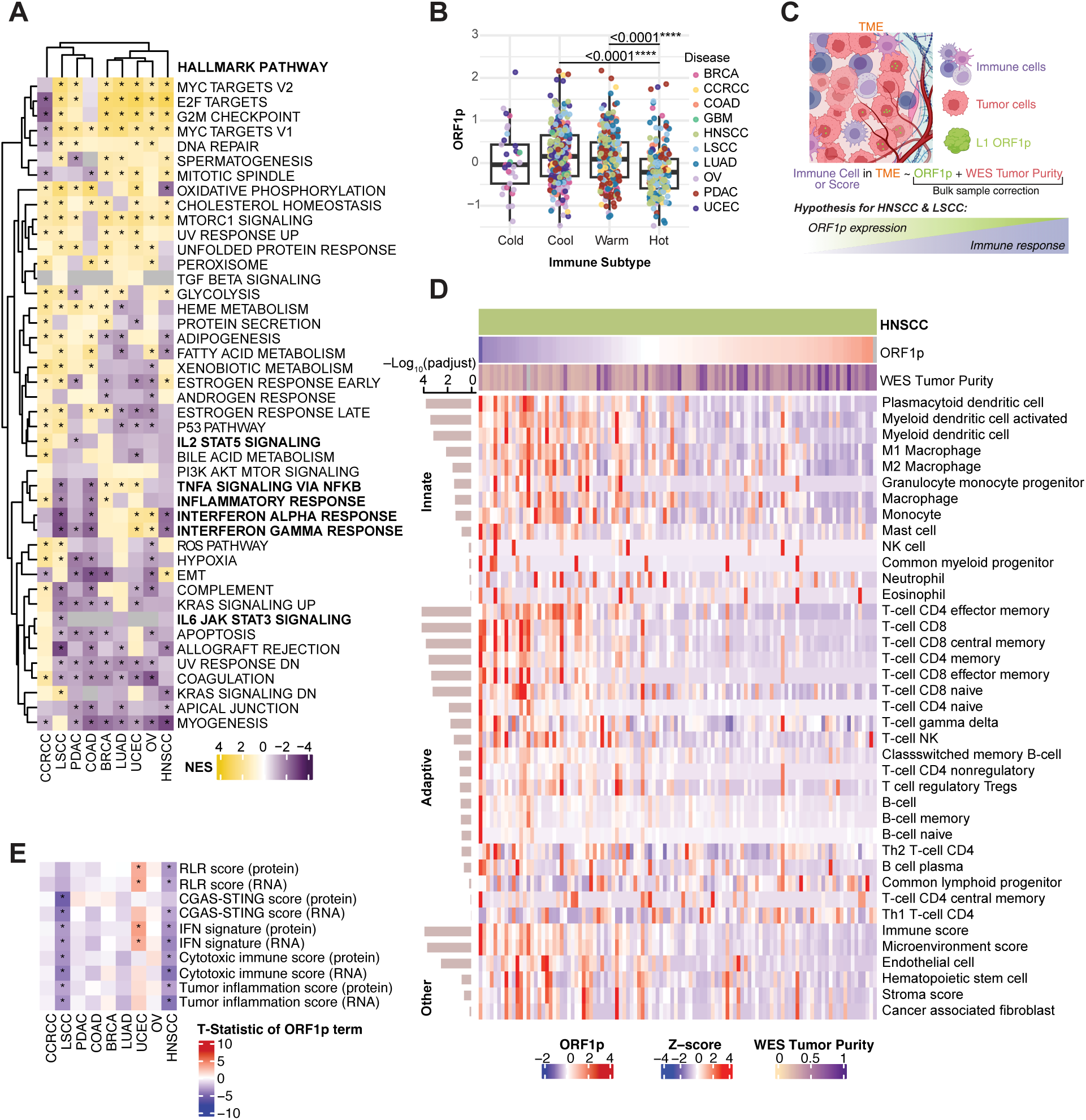
ORF1p expression regulates immune pathways and immune cell infiltration in a tumor-specific manner. (A) Heatmap of normalized enrichment scores (NES) from GSEA of HALLMARK pathways. Input is a gene list ranked by decreasing logFC from differential proteomic expression analysis (per cancer type). Rows are pathways and columns are cancer types. Asterisk (*) indicates significant NES (adjusted P < 0.01). Immune-related pathways are bolded. (B) ORF1p abundance by CPTAC immune subtypes from Li et al., 2023^77^: Cold, Cool, Warm and Hot. Each point is a sample colored by cancer type. Two-sided Wilcoxon rank sum test (BH-adjusted P values). (C) Schematic of tumor microenvironment (TME) explaining linear regression model used in panels D-E and decreased immune response in ORF1p-high squamous cell carcinoma tumors. (D) Heatmap of xCell immune cell type deconvolution scores ordered by increasing ORF1p expression in HNSCC. WES tumor purity covariate also annotated. Rows are immune cell types grouped by branches of the immune system. Rows within each group are ordered by decreasing significance of the P adjusted value of the ORF1p term from the linear regression model. (E) Heatmap of T-statistic from linear regression predicting the immune score (row) by ORF1p expression and tumor purity. Scores are calculated based on RNA and protein expression. Asterisk (*) indicates significant association with ORF1p expression (P adjusted < 0.01). Positive T-statistic indicates positive correlation, while negative T-statistic indicates negative correlation. For box plots: p ≤ 0.05, **p≤ 0.01, ***p ≤ 0.001, ****p ≤ 0.0001.

These data suggest that, in specific cancer types, tumors with higher expression of ORF1p downregulate inflammatory pathways with potential implications in tumor immune response. To explore this hypothesis we further categorized tumors into immune subtypes (cold, cool, warm and hot) based on a previously published CPTAC pan-cancer study linking oncogenic drivers to functional states (Figures 4B and S4A-I).^77^ When comparing ORF1p expression by immune subtype at the pan-cancer level, we observed significantly lower ORF1p expression in immune-hot tumors compared to those in the immune-warm and immune-cool subtypes (Figure 4B). Similar significant trends were observed at the individual cancer level for COAD, HNSCC, LSCC and LUAD (Figure S4C-F), validating pathway-based clustering (Figure 4A). Assessing L1 RNA expression by immune subtypes, we also observed significantly lower L1 RNA in the immune-hot subtype at the pan-cancer level and individually for HNSCC, LSCC and LUAD (Figure S4D-F). Together, these findings suggest a negative relationship between L1 expression and host immune response in the squamous cell carcinomas (Figure 4C).

To further evaluate this relationship, we performed regression analyses to predict the effect of ORF1p expression on immune cell type. RNA-derived xCell^78^ immune cell type estimations were used as model input, correcting for WES-based tumor purity for bulk sample correction (Figure 4C). The only statistically significant finding at the pan-cancer level was a negative correlation between ORF1p expression and activated myeloid dendritic cells (Figure S5A). Independent, cancer-specific regression revealed that the associations between ORF1p expression and immune cell infiltration are unique to each tumor type (Figures 4D and S5B-I). Significant associations (adjusted *P* < 0.05) between ORF1p expression and immune cell activation were only identified in HNSCC, LSCC and LUAD. The top significantly associated cell types with ORF1p expression in HNSCC and LSCC tumors, were, respectively, plasmacytoid dendritic cells (pDCs) and activated myeloid DCs in the innate arm and CD4+ effector memory T-cells and class-switched memory B-cells for the adaptive arm (Figures 4D and S5F). The involvement of pDCs is particularly notable given their role as key regulators of antiviral immunity via inflammatory cytokine and their ability to differentiate into myeloid DCs^79^, that we also found negatively correlated with ORF1p expression in HNSCC (adjusted *P* < 0.05) (Figure 4D).

This data corroborate our hypothesis that ORF1p expression coincides with reduced innate immune response, potentially compromising tumor immunosurveillance.^80,81^ This findings are also in line with our pathway-level expression analysis of the interferon alpha and gamma response and the MAPK/NF-𝜿B axis (TNF-a signaling via NF-𝜿B) that shows depletion of these pathways in ORF1p-high tumors of both HNSCC and LSCC (Figure 4A and Table S3). These pathways are indeed known to drive pDC maturation into activated myeloid DCs^82,83^ and their downregulation suggests that high L1 ORF1p expression is associated with a microenvironment depleted of proinflammatory signaling resulting in fewer functional DCs. In contrast, molecular and pathway analysis in UCEC show opposite trends compared to HNSCC and LSCC (Figures 2B-J, 3D and 4A). Notably in UCEC, ORF1p expression has a positive correlation with immune activation (adjusted *P* value < 0.1), with the strongest association observed for activated myeloid DCs (Figure S5I). These results show that ORF1p overexpression has tumor-type specific effects on immune infiltration, dividing CPTAC cancers into two distinct categories: those where ORF1p expression correlates with immune suppression (e.g. HNSCC and LSCC) and those where it correlates with activated immune response (e.g., UCEC). This dichotomy is consistent with previously conflicting reports of both positive and negative immune correlations with L1 activation.^22,29,30,42^

To gain insight into the molecular mechanisms that drive the functional interaction between L1 ORF1p expression and the immune response, we derived a series of RNA- and protein-based immune scores representing cytosolic dsRNA triggered responses by RIG-I-like receptors (RLR score) and cytosolic dsDNA or DNA:RNA hybrids triggered pathways (cGAS/STING scores). We also included IFN signaling, cytolytic activity of the adaptive immune response, and tumor inflammation scores based on previous published signatures (Methods). We built cancer-specific linear regression models to predict each immune score by ORF1p expression. Corroborating our previous analyses, we found that HNSCC and LSCC exhibited similar response to ORF1p expression, with clear negative correlations between ORF1p expression and nearly all RNA- and protein-based immune scores (Figure 4E, adjusted *P* value < 0.01). Notably, and again consistent with our previous findings, UCEC showed an opposite association between ORF1p and immune response, with positive associations identified between ORF1p expression and RLR and IFN signature scores (Figure 4E). These findings also suggest that in UCEC repeat element-derived molecular patterns likely trigger mostly an antiviral response through dsRNA sensing mediated by RIG-I and/or MDA5 rather than cytosolic dsDNA or DNA:RNA hybrid detection triggered by the cGAS/STING pathway.

### ORF1p expression predicts cancer-specific aneuploidies

To investigate the underlying reasons why the immunogenicity of ORF1p overexpression varies across cancer types, we evaluated the relationship between somatic copy number alterations (SCNAs) and ORF1p. Aneuploidy, the state of having gained or lost chromosomes, is a fundamental hallmark of cancer.^84^ Aneuploid states in specific cancer types have been linked to immune evasion and altered immune cell infiltration.^85^ We argue that if tumors attempt to decrease inflammation and IFN signaling for their growth and development, a potential solution to achieve this goal would be the loss or gain of genomic regions rich in genes involved in immune regulation (e.g. the IFN cluster on chromosome 9p).^86^ We therefore assessed the correlation of chromosome arm-level copy number (CN) ratios with ORF1p expression and found cancer-specific significant associations (adjusted *P* value < 0.05) (Figure S6A). Interestingly, as with our previous analyses, HNSCC and LSCC followed similar trends in the arm-level aneuploidies correlating with ORF1p expression, with chromosomes 1q gain, 2p gain, 3q gain, and 4p loss significantly correlating with ORF1p expression in both tumor types (Figure S6A).

We next binarized cytoband-level copy number ratios as gains (CN > 0.3) or losses (CN < -0.3) and performed individual generalized regressions to determine whether ORF1p expression can predict the gain or loss of specific cytogenic bands across cancer types (Methods). We found that ORF1p significantly (adjusted *P* < 0.05) predicts 1q gain in both HNSCC and LSCC (Figures 5A and S6B). Chromosome 1 is a chromosome known to contain several immune regulating genes.^87^ When assessing binary chromosome 1q gain status in HNSCC and LSCC, tumors with 1q gain in both cancer types had significantly higher ORF1p expression (Figures 5B and S6C). In UCEC, ORF1p expression was correlated with different significant gains and losses (Figure 5C), chief among them gain of chromosome 2p. Indeed, UCEC tumors with 2p gain have significantly higher ORF1p expression than those with no amplification (Figure 5D)

**Figure 5.**
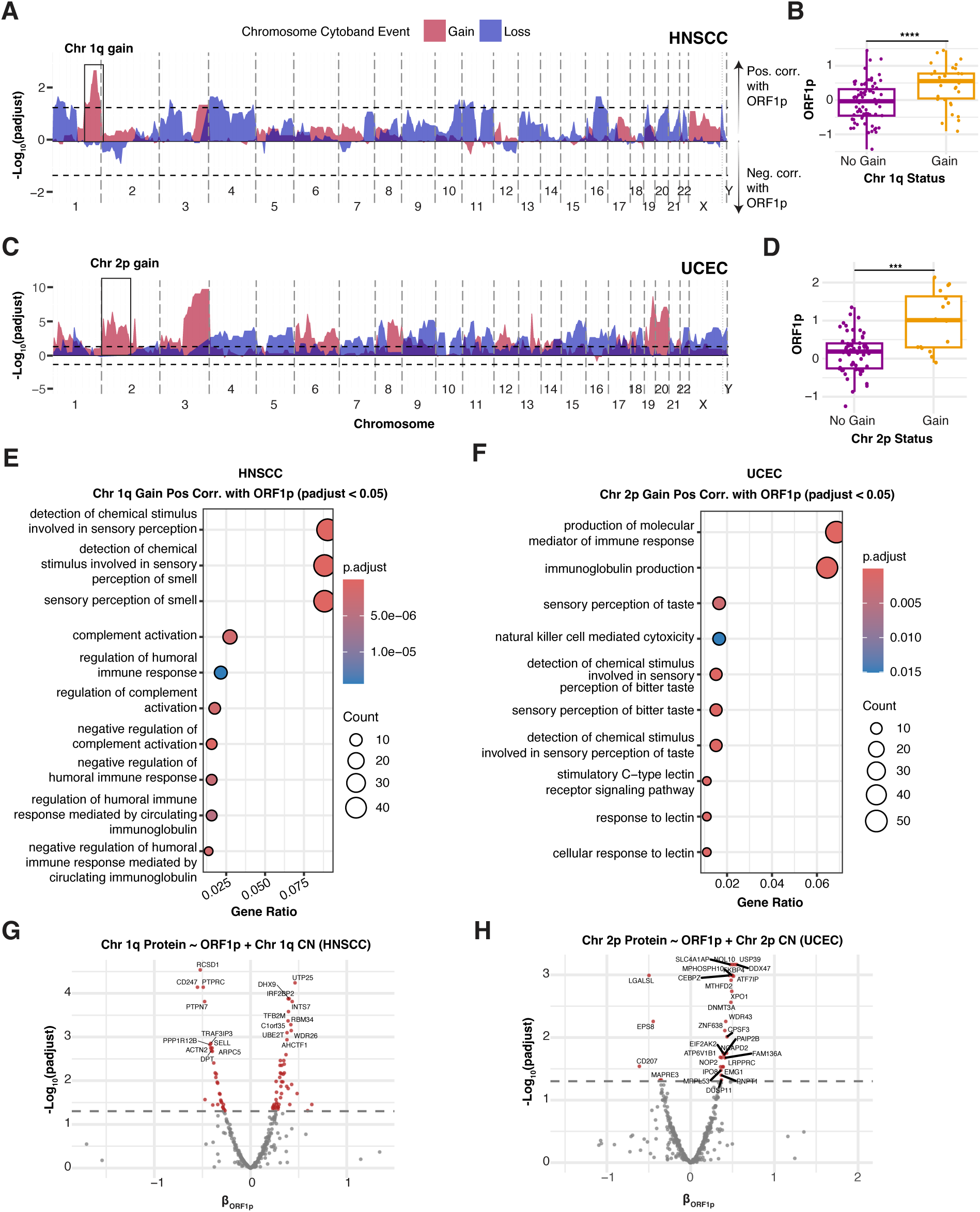
ORF1p expression predicts cancer-specific aneuploidies. (A) Area plot represents the association between chromosome cytoband gains (red) and losses (blue) and ORF1p expression in HNSCC in a multivariable logistic regression model (see Methods for formula). Each chromosome is split by a grey vertical dotted line. The black horizontal dotted line represents adjusted P value = 0.05. Positive - Log10(adjustedP) indicates the cytoband gain or loss is positively associated with ORF1p expression, while a negative -Log10(adjustedP) indicates the cytoband gain or loss is negatively associated with ORF1p expression. Chromosome 1q gain is boxed to emphasize further analysis in B, E, and G. (B) ORF1p expression in HNSCC is stratified by gain or no gain of 1q. (C) Area plot represents the association between chromosome cytoband gains (red) and losses (blue) and ORF1p expression in UCEC. Chromosome 2p gain is boxed to emphasize further analysis in D, F and H. (D) ORF1p expression in UCEC is stratified by gain or no gain of 2p. (E) Dot plot showing top 10 GOBP pathways from ORA for ORF1p-predicted (adjusted P < 0.05) gained 1q cytobands that are positively associated with ORF1p in HNSCC. (F) Dot plot showing top 10 GOBP pathways from ORA for ORF1p-predicted (adjusted P < 0.05) gained 2p cytobands that are positively associated with ORF1p in UCEC. (G) Volcano plot -Log10(adjustedP) vs. β-coefficient of ORF1p from linear model predicting protein expression of genes located on significantly predicted (adjusted P < 0.05) gained 1q cytobands in HNSCC. Arm-level CN ratio (continuous) is used as a covariate to determine associations independent of CN. (H) Volcano plot -Log10(adjustedP) vs. β-coefficient of ORF1p from linear model predicting protein expression of genes located on significantly predicted (adjusted P < 0.05) gained 2p cytobands in UCEC.

Given the strong prediction of ORF1p for unique CN alterations in HNSCC, LSCC and UCEC, we investigated whether specific CN alterations in tumors might enable high ORF1p expression through the amplification or loss of specific pathways. We categorized CN alterations into 4 groups: those (1) positively associated with ORF1p and CN gain (Pos-Gain); (2) positively associated with ORF1p and CN loss (Pos-Loss); (3) negatively associated with ORF1p and CN gain (Neg-Gain); and (4) negatively associated with ORF1p and CN loss (Neg-Loss) and filtered for an adjusted *P* < 0.2 (Figure S6D). We then performed over-representation analysis (ORA) on each of these gene sets per cancer type, determining which biological pathways were enriched for ORF1p-predicted aneuploidies grouped into each category (Table S4). Next, we performed similar ORA for genes on the specific chromosome arms significantly predicted by ORF1p in HNSCC (1q gain), LSCC (1q gain) and UCEC (2p gain) to determine pathways unique to each arm (Tables S5).

As expected, we detected a significant enrichment for negative regulation of humoral immune response and negative regulation of immune effector processes, for genes on chromosome 1q in HNSCC and LSCC (Figure 5E and S6E). Conversely, we identified pathways involved in immunoglobulin production and natural killer cell-mediated cytotoxicity for genes enriched on chromosome 2p in UCEC (Figure 5F). Together, these findings suggest that ORF1p expression is associated with aneuploidies enriched in genes involved in immune response and that aneuploidies found in HNSCC and LSCC versus UCEC can explain the ORF1p immune dichotomy.

To better understand the protein-level impacts of ORF1p-associated aneuploidies in HNSCC and UCEC, we isolated genes located on specific 1q and 2p cytoband regions significantly (adjusted *P* < 0.05) predicted by ORF1p expression using overall 1q and 2p CN ratios as covariates, respectively, to identify expression relationships independent from total copy number dosage (Figures 5G-H). In both HNSCC and UCEC, we identified proteins negatively correlated with ORF1p despite overall amplification of their respective chromosomes, indicating that chromosome copy number does not by itself explain proteomic associations with ORF1p for these genes and that additional ORF1p- dependent mechanisms may contribute to their deregulation.

Altogether, these findings suggest that ORF1p expression can direct the formation of specific aneuploidies that modulate innate immune and immune cell activation pathways. Further experimental validation is needed to determine whether derepression of TEs in early tumorigenesis leads to L1/IFN-permissive CN alterations or if ORF1p expression and specific CN alteration develop independently through a common causative event.

## DISCUSSION

To our knowledge, this work provides the first large-scale proteogenomic pan-cancer study of L1 with patient-level L1 insertion, RNA and ORF1p quantifications and systematic integration with multi-omic data types. By leveraging CPTAC pan-cancer proteogenomic data, we demonstrate that ORF1p-high tumors are characterized by unique proteogenomic and immune changes that are tumor-specific and related to copy number alterations, potentially resulting in a permissive molecular environment for L1 expression (Figure 6).

**Figure 6.**
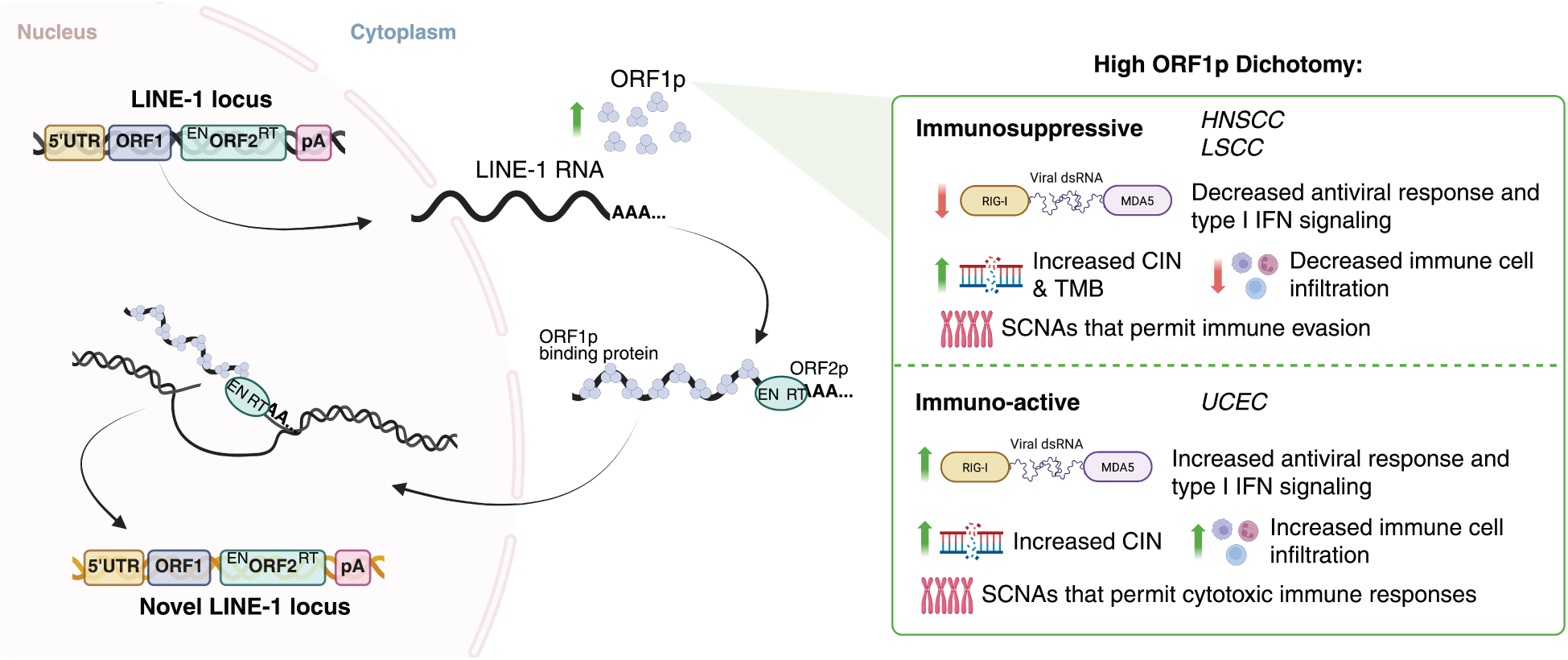
Summary schematic of ORF1p-high dichotomy. ORF1p-high tumors can be divided into two groups: an (1) immunosuppressive group characterized by negative correlations between ORF1p expression and antiviral responses and innate immune signaling, increased CIN and TMB, and decreased immune cell infiltration in the background of copy number alterations that permit immune evasion, and an (2) immuno-active group characterized positive correlations between ORF1p expression and antiviral/innate immune responses, increased CIN and immune cell infiltration in the background of copy number alterations that permit cytotoxic immune functions.

Our comprehensive analysis reveals that ORF1p expression and its molecular consequences subdivide tumors into two distinct groups (Figure 6). The first is an immunosuppressed group, including HNSCC and LSCC, which exhibited high ORF1p expression, especially in classical subtypes, and negative associations between ORF1p expression and immune scores and cell types. Consistent with previous studies showing particularly high L1 permissiveness in squamous cell carcinomas, heightened ORF1p expression in HNSCC and LSCC may be linked to the high prevalence of p53 mutations in these tumors, as p53 is a direct repressor of L1.^33,34^ Within these cancer types the highest ORF1p levels were observed in the classical subtypes, both of which have been associated with high chromosomal instability, strong smoking evidence and p53 mutation in each respective tumor type.^60^ Given that current and past smokers with HNSCC, irrespective of molecular subtype, had significantly higher ORF1p expression than non-smokers, our findings corroborate existing evidence that hypomethylation of L1 is associated with smoke-exposed epithelia and that this hypomethylation is associated with an increased risk of HNSCC.^88^ These squamous cell carcinomas also show negative correlations between ORF1p expression and immune infiltration, particularly plasmacytoid and activated myeloid dendritic cells. The association between ORF1p expression and 1q gain in HNSCC underscores the potential role of ORF1p overexpression in reshaping immune-regulatory circuits through genomic alterations. Not only are the observed over representation of genes on 1q linked to the negative regulation of immune processes^87^, but ORF1p is negatively correlated with proteins located on 1q independent of 1q copy number dosage. The top 3 most significantly anticorrelated 1q proteins with ORF1p included RCSD1, a cytoskeletal regulatory molecule that has been shown to associate with immune infiltrating cells^89^, CD247, the gene encoding the essential CD3 zeta (ζ) chain component of the T cell receptor (TCR) complex^90^, and PTPRC (or CD45), a hematopoietic-specific transmembrane protein tyrosine phosphatase that regulates Src kinases required for T- and B-cell antigen receptor signal transduction.^91^ Though further *in vitro* and *in vivo* experiments are needed to determine whether ORF1p is a cause or consequence of SCNAs, our findings suggest that copy number alone does not explain the association of immunosuppressive characteristics with ORF1p expression in these tumor types.

Conversely, the second group of tumor types is an immune-active group, including UCEC tumors, that show the opposite trend, with positive associations between ORF1p expression and immune cell infiltration, particularly activated myeloid DCs and upregulation of antiviral response pathways. UCEC tumors with the highest ORF1p expression were identified in the CNV-H subtype known for high copy number variations, genomic instability and elevated expression of cell cycle and antiviral response proteins.^61^ In this tumor type, one of the most significant copy number alterations predicted by ORF1p expression were gains in chromosome 2p. Unlike 1q, over-representation analysis in chromosomal region 2p is linked to pathways that foster immune activation, such as immunoglobulin production and natural killer cell-mediated cytotoxicity, suggesting a distinct mechanism of immune modulation in UCEC compared to the squamous cell carcinomas. Independent of chromosome 2p amplification, 2p proteins were still significantly positively correlated with ORF1p, including antiviral dsRNA sensor EIF2AK2 (PKR)^92^ and USP39, a deubiquitinase that stabilizes STAT1 to sustain type I IFN immunity.^93^ This disparity raises intriguing questions about the role of ORF1p overexpression on shaping cancer-specific aneuploidies and immune responses, which could be reflective of the unique evolutionary pressures or tissue-specific contexts inherent to each cancer^94^, for example squamous cell carcinomas (HNSCC and LSCC) versus adenocarcinomas (UCEC) in our study.

This dichotomy suggests cancer specific adaptations to L1 derepression and our data indicate that ORF1p expression may compromise tumor immunosurveillance in squamous cell carcinomas by reducing type I IFN activation and subsequent pDC infiltration. Further, our data suggest that reduced infiltration of key immune cells, including plasmacytoid and activated myeloid dendritic cells in HNSCC and LSCC, potentially compromises tumor immunosurveillance by decreasing type I interferon secretion and adaptive immune response recognition in these tumor types. Further experimental validation is needed to connect ORF1p’s role in masking dsRNA from repetitive elements and precise mechanisms by which ORF1p may modulate immune response in the context of immune cell type recruitment.^2942^

Our study also offers valuable insights into the role of ORF1p expression in modulating global protein and phosphoprotein expression. We identified PPM1G phosphatase upregulation in ORF1p high tumors as particularly compelling, as ORF1p- high HNSCC tumors upregulate PPM1G to negatively modulate immune activation by regulating post-translational modification of innate immune adapter proteins and subsequent type I IFN gene expression programs, similar to the immune evasion effect caused by hijacking of PPM1G during Kaposi’s sarcoma-associated herpesvirus (KSHV) infection.^69^ The observed reduction in STING1 and MAVS phosphorylation in ORF1p high HNSCC tumors suggests that PPM1G overexpression dampens antiviral responses, facilitating a tolerogenic environment that supports L1 overexpression. Additionally, proteins upregulated in ORF1p-high tumors include those in RNA and RNP metabolism, as ORF1p-high tumors may upregulate these proteins to stabilize L1 RNA expression and support active L1 translocation.^9,67,95^

This study improves our understanding of ORF1p overexpression in cancer, but several questions remain. First, though we identify significant associations of ORF1p with proteins, immune pathways and aneuploidies, precise molecular mechanisms remain unclear and need more basic experimental follow-up. Second, sample sizes for specific demographic variables and somatic mutation statuses, such as p53, varied per cancer type, warranting validation analyses with independent proteogenomic datasets. In the age of personalized medicine, our findings advance ORF1p as a candidate biomarker by introducing novel tumor-specific associations of ORF1p with immune infiltration and copy number alterations that can enhance clinical utility.^75,96–98^ Emerging development of ultrasensitive immunoassays for detection of plasma-circulating ORF1p can be leveraged to predict immune checkpoint inhibitor (ICI) resistance, especially in ORF1p-high tumors, and to longitudinally monitor immune adaptation to intercept emerging resistance during ICI therapy.^99–101^

Our comprehensive pan-cancer characterization of L1 ORF1p provides valuable insights into the molecular underpinnings of L1 activity in cancer. These findings pave the way for future research to explore the therapeutic potential of targeting ORF1p and its associated molecular consequences in specific cancer treatments.

## METHODS

### CPTAC Pan-Cancer Data

All data used in this study can be downloaded from the NCI’s Proteomic Data Commons (PDC) at https://proteomic.datacommons.cancer.gov or https://proteomic.datacommons.cancer.gov/pdc/cptac-pancancer. Details regarding sample acquisition, harmonization of the pan-cancer data processing, demographic composition and methods can be found in the CPTAC pan-cancer resource paper.^43^

### Human Subjects

In this study, we used proteogenomic and clinical data obtained by the Clinical Proteomic Tumor Analysis Consortium (CPTAC) from over 1,000 eligible participants. This cohort (n = 1,072) spans 10 cancer types: breast adenocarcinoma (BRCA, n = 122)^51^, clear cell renal cell carcinoma (CCRCC, n = 110)^56^, colon adenocarcinoma (COAD, n = 96)^52^, glioblastoma multiforme (GBM, n = 99)^102^, head and neck squamous cell carcinoma (HNSCC, n = 110)^55^, lung squamous cell carcinoma (LSCC, n = 108)^58^, lung adenocarcinoma (LUAD, n = 110)^57^, pancreatic ductal adenocarcinoma (PDAC, n = 140)^59^, high-grade serous ovarian cancer (OV, n = 82)^53^ and uterine corpus endometrial carcinoma (UCEC, n = 95).^54^ Protocols and consent documentation were reviewed by the institutional review boards at the tissue source sites, ensuring compliance with CPTAC guidelines. All specimens were collected from patients during routine tumor resections, with informed consent obtained from the patients, in accordance with CPTAC standards and guidelines.

### Clinical Data

Clinical information was downloaded from CPTAC Data Coordinating Center (DCC). Clinical variable names and units were standardized and unified. Clinical and molecular data can be accessed and downloaded from the NCI’s Proteomic Data Commons (PDC) at https://proteomic.datacommons.cancer.gov.

### Specimen Acquisition

CPTAC collected tumor and normal adjacent tissue (NAT) and whole blood samples derived from patients diagnosed with BRCA, CCRCC, COAD, GBM, HNSCC, LSCC, LUAD, OV, PDAC and UCEC. All cases in the study met the histology acceptance criteria, and detailed information regarding sample processing can be found in each respective flagship publication and the CPTAC pan-cancer resource paper.^43^

### Somatic Copy Number Variation (CNV) Data

CNV ratios were derived from whole genome (WGS) and whole exome sequencing (WES) for each tumor type as described in Li et al., 2023.^43^

### Somatic Mutation Data, Mutational Burden and Signatures

Somatic mutation calls from WES data for matched tumor/normal samples were analyzed using a hg38 WES characterization pipeline and harmonized as described in Li et al., 2023.^43^ Estimated tumor mutational burden and mutational signatures were derived using the Genomic Identification of Significant Targets in Cancer (GISTIC2.0)^103^ algorithm and MutSig2V^104^ as described in Li et al., 2023.^43^

### RNA Expression Data

Global RNA expression data was obtained from gene-level read counts (Fragments Per Kilobase of transcript per Million mapped reads, FPKM) and processed as described in Li et al., 2023.^43^ Data can be downloaded from the Genomic Data Commons (GDC) at https://portal.gdc.cancer.gov/ or from https://proteomic.datacommons.cancer.gov/pdc/cptac-pancancer.

### Cell Type Enrichment Deconvolution (RNA)

Cell type abundance was inferred by the xCell web tool^78^, which performed the cell type enrichment analysis from gene expression data for 64 immune and stromal cell types (default xCell signature) as described in Li et al., 2023.^43^ The xCell immune score per sample that integrates the enrichment scores B cells, CD4+ T-cells, CD8+ T-cells, DC, eosinophils, macrophages, monocytes, mast cells, neutrophils, and NK cells. The xCell microenvironment score is the sum of the immune score and stroma score.

### Protein Expression Data

Global protein expression data and peptide-spectrum match (PSM) counts were processed as described in Li et al., 2023^43^ and can be downloaded from the PDC.

### Phosphoprotein Expression Data

Global, site-level phosphoprotein expression data from phosphopeptide-enriched proteomic data were processed as described in the methods of Li et al., 2023.^43^ Data can be downloaded from the PDC.

### Adjustment of Phosphoprotein Abundance by Protein Expression

To analyze phosphorylation changes that happen specifically at the phosphoprotein level, we performed a linear regression between phosphosite abundance (BCM pipeline) and protein abundance (BCM pipeline) of each phosphosite using the base R package stats (lm function). We then computed the response residuals for each phosphosite, which normalizes the difference between the observed protein abundance and expected value based on protein expression (residual) by their standard deviation.

### Adjustment of Somatic Copy Number by Tumor Purity and Ploidy

Gene-level copy number and chromosome arm-level copy number tables were adjusted for tumor purity and ploidy. The log2 copy number is first linearized. The normal cell contribution to the copy number signal is calculated by multiplying the fraction of normal cells in the sample by the deviation of the sample from diploid (normal cell is assumed to be diploid) and by the integer 2. The normal cell contribution is then divided by the tumor purity multiplied by the ploidy to determine the correction which is then added back to the linearized copy number input and log2 transformed. This adjustment corrects the tumor copy number by unmixing the tumor signal from normal cell contamination and adjusts for a non-diploid tumor baseline.

### Calculation of Chromosome Arm and Cytoband-Level Copy Number

WES log2 gene-level CNV matrices from the BCM pipeline were used to derive chromosome arm-level and cytoband-level copy number matrices per cancer type by calculating the mean copy number of all genes for that chromosome arm or cytoband.

### Calculation of Aneuploidy Score

To estimate the extent of tumor aneuploidy, we calculated the sum of the absolute value of the copy number for each chromosome arm per cancer type.

### L1 Quantifications

#### Intact L1 RNA

L1 RNA was quantified as described in McKerrow et al., PNAS, 2022^44^ by passing available RNA sequencing BAM files from the GDC to L1EM^105^ on the Cancer Genomics Cloud. L1EM utilizes expectation maximization to estimate locus-specific L1 expression and to separate proper L1 expression from passive co-transcription that includes L1 RNA but does not support retrotransposition. For intact L1 RNA quantifications, full length L1 loci with no stop codon in either ORF1 or ORF2 were considered expressed if at least two read pairs per million (FPM) were assigned to that locus and less than 10% of the RNA assigned to that locus was estimated to be passive co- transcription. Total intact L1 RNA expression was estimated by adding together the FPM values for each such locus. This number is an estimate of the total amount of active L1 RNA expression from intact loci. By “active” we mean an expression that is driven by the L1 promoter (i.e. the TSS is within 25 bp of the start of L1). Samples were excluded if no locus was detected at 2 FPM as this may be due to data quality rather than a lack of L1 RNA.

### L1 ORF1p

L1 ORF1p was quantified as described in McKerrow et al., PNAS, 2022.^44^ ORF1p was quantified from tandem mass tag (TMT) labeled mass spectrometry (MS/MS) proteomics data. X!Tandem^106^ was used to search mass spectra against a combined database that included both the standard Ensembl human proteome and in silico translated L1 ORF1 sequences from hg38 to the human Ensembl proteome. Oxidation of methionine (+15.994915@M) was included as a potential modification. Carbamidomethylation of cysteine (+57.022@C) was set as a fixed modification in addition to modifications appropriate to the particular isobaric labeling used (+144.102063@[, +144.102063@K for iTRAQ4 and +229. 162932@[, +229.162932@K for TMT10). Peptides were quantified by calculating the log ratio between the reporter intensity for each sample and the reporter intensity of an internal control. Peptide/peptide spearman correlations were calculated and peptides that had a correlation of at least 0.6 with two other peptides were retained. We then identified matches to 20 ORF1p peptides: DFVTTRPALK, EALNMER, EWGPIFNILK, LIGVPESDGENGTK, LIGVPESDVENGTK, LSFISEGEIK, LTADLSAETLQAR, NLEECITR, NVQIQEIQR, QANVQIQEIQR, REWGPIFNILK, RNEQSLQEIWDYVK, SNYSELR, SNYSELREDIQTK, VSAMEDEMNEMK, YQPLQNHAK, DFVTTRPALQELLK, LENTLQDIIQENFPNLAR, VSAMEDEMNEMKR, and NEQSLQEIWDYVK. If a peptide were detected multiple times in the same sample, the median across peptide spectral matches was used. Then to get a protein quantification, the median was taken across all peptides in the preceding list that were identified in that sample. Finally, the quantification was translated by the median log ratio for all human proteins to account for variation in the size of the input sample. This pipeline was implemented and executed as a workflow on the Cancer Genomics Cloud.

### High Confidence Somatic Insertions

Somatic insertions were identified as described in McKerrow et al., PNAS, 2022^44^ by using the Mobile Element Locator Tool (MELT)^107^ v2.1.15 on the Cancer Genomics Cloud. MELT was run on each pair of matched cancer/normal WGS datasets (no WGS for BRCA, OV, and COAD). Only insertions with the greatest evidence (ASSESS = 5) were considered. To be considered a high-confidence somatic insertion, the insertion had to be called heterozygous in cancer and homozygous absent in normal, with the log likelihood of both of these genotypes being at least 10 greater than the log likelihood of the next most probable genotype.

### ORF1p-High versus ORF1p-Low Analyses

#### Derivation of ORF1p-High and ORF1p-Low Groups

ORF1p-high and OR1p-low groups were derived using quartiles from the distribution of ORF1p expression per individual cancer type. Samples whose ORF1p expression was less than or equal to the first quartile were considered “low”, and samples whose ORF1p expression was greater than or equal to the third quartile were considered “high”. Samples not labeled as “high” or “low” were excluded from differential expression analyses.

### Differential Proteomic Expression Analysis

The DEqMS R package^108^ was used to perform differential protein expression analysis using proteomic data for each individual cancer type comparing ORF1p-high versus ORF1p-low samples. DEqMS is a robust method for both labelled and label-free MS data that considers peptide spectrum matches (PSMs) per protein to adjust for the inherent dependence of protein variance on the number of peptides used for quantification. The consideration for the actual MS-proteomics data structure improves the accuracy of differentially expressed protein (DEP) detection. Individual observed and normalized log2 reference intensity protein abundance tables from the BCM pipeline were used as input for each cancer type, filtered for ORF1p-high and ORF1p-low samples, along with matched PSMs obtained from the UMich-GENCODE34 proteomic tables. ORF1p expression was not included in these matrices, and rows with NAs were removed. WES tumor purity was used as a covariate in the model. For samples with missing tumor purity values, we used the mean tumor purity of all samples in the analysis.

### Gene Set Enrichment Analysis for Differential Protein Expression

The clusterProfiler R package^109^ was used to perform GSEA for human MSigDB hallmark gene sets. Proteins ordered by decreasing logFC from DEqMS were used as the gene list input. The minGSSize was set to 30, the maxGSSize was set to 500, and the pvalueCutoff was set to 1 to capture all results. The pAdjustMethod was set to “BH”.

### RNA- and Protein Expression-Based Scores

#### Calculation of Tumor Inflammation Score

The tumor inflammation signature is a published set of 18 genes, including CD276, HLA- DQA1, CD274, IDO1, HLA-DRB1, HLA-E, CMKLR1, PDCD1LG2, PSMB10, LAG3, CXCL9, STAT1, CD8A, CCL5, NKG7, TIGIT, CD27, and CXCR6, that enrich for clinical benefit to immune checkpoint blockade by measuring the level of tumor microenvironment inflammation.^110^ The score was calculated as referenced by ranking the gene expression values within each sample and taking the weighted average of the gene ranks within each sample. RNA and protein expression was used to infer tumor inflammation score, however RNA-based scores had less gene missingness.

### Calculation of Cytotoxic Immune Score

The cytotoxic immune score was calculated as referenced in Davoli et al., Science, 2017^85^ by ranking the expression of each gene per sample and then taking the sum of the ranks per sample. The gene set captures the expression of genes characteristic of the adaptive immune response and immune-mediated cytolytic and cytotoxic activities and includes CD247, CD2, CD3E, GZMH, NKG7, PRF1, and GZMK. RNA and protein expression was used to infer cytotoxic immune score.

### Calculation of CGAS-STING, RLR, and Interferon Signature Scores

These scores were inferred by single sample scoring using the singscore method from the singscore R package.^111^ This method does not depend on background samples resulting in scores that are stable regardless of the composition and number of samples being scored. The CGAS-STING activation score (cytosolic dsDNA sensing score) includes CGAS, STING1, CXCL10, IRF3, TBK1, and STAT1.329,357 The RLR score (cytosolic dsRNA sensing score) includes DDX58 (RIG-I), IFIH1 (MDA5), MAVS, EIF2AK2 (PKR), and IRF3.^42,92,112^ The curated list of 140 IFN and ISGs from Sun et al., Immunity, 2024^29^ was used as the IFN signature score. RNA and protein expression was used to infer all scores.

### Survival Analysis

The survival^113^ and survminer^114^ R packages were used to perform survival analysis. The Kaplan-Meier curve of overall survival was used to compare the pan-cancer prognosis among ORF1p status. Log-rank test (survminer) was used to test the differential survival outcomes between ORF1p-high and ORF1p-low groups.

### Correlations and Linear Regressions

Correlations were run using the cor.test() function in R. The method argument was set to “spearman” and the alternative argument set to “two.sided”. Linear regressions were run to account for covariates, such as tumor purity or CN ratio, using the R functions lm() for predicting continuous variables and glm() to predict binary (0, 1) outcomes, such as the prediction of chromosome gain (0, 1) or loss (0, 1).

### Over-Representation Analysis (ORA) of Chromosomal Aneuploidies

#### ORA for All Gained or Lost Chromosomes Associated with ORF1p Expression

Individual linear models were run per cancer type to predict the binary (0,1) gain (CN > 0.3) or loss (CN < -0.3), as previously described, of cytogenic bands by ORF1p expression using tumor purity as a covariate:

1. *Gain (1) or No Gain (0) ∼ ORF1p + WES Tumor Purity*
2. *Loss (1) or No Loss (0) ∼ ORF1p + WES Tumor Purity*

Results were filtered for ORF1p associations with adjusted P values < 0.2. Cytogenic bands were stratified based on the direction of ORF1p association (positive or negative) and type of copy number event (gain or loss). Four gene lists, Pos-Gain, Pos-Loss, Neg- Gain and Neg-Loss were constructed from genes within all cytogenic bands that fit each respective category and used as input for ORA to determine overall pathways genomically gained or lost.

### ORA for Individual Gained or Lost Chromosomes Associated with ORF1p Expression

To determine pathways unique to specific chromosomal aneuploidies associated with ORF1p, we performed similar ORAs as described above for individual chromosome arm aneuploidies. This time, results were more strictly filtered for significance (adjusted P < 0.05), and genes located on individual chromosome arms (p or q) were aggregated into gene lists. ORA was run for each individual significant aneuploidy to determine which pathways enriched on unique chromosome arms are associate with ORF1p expression.

### Gene Set Collections for ORA

Gene set collections were obtained from the Molecular Signatures Database (MSigDB) using the msigdbr R package.^115^ ORA was performed using multiple complementary approaches to capture diverse biological processes and pathways:

Gene Ontology Biological Process (GO:BP) ORA was assessed using the enrichGO function from the clusterProfiler R package.^109^ Analysis parameters included a minimum gene set size of 10, maximum gene set size of 500, and BH multiple testing correction. The background universe consisted of all genes present in the regression analysis. Hallmark Pathway ORA was performed using the enricher function with MSigDB Hallmark gene sets as the reference database. The same statistical parameters and background universe were applied as for GO:BP analysis. BioCarta Pathway ORA was evaluated using BioCarta gene sets with identical statistical parameters to ensure consistency across analyses. For KEGG Pathway ORA, gene symbols were converted to Entrez IDs using the bitr function from the org.Hs.eg.db annotation package.^116^ Enrichment was performed using enrichKEGG function with organism set to "hsa" (Homo sapiens). Reactome Pathway ORA was conducted similar to KEGG analysis using the enrichPathway function with Entrez IDs and organism set to "human". Gene lists containing fewer than 5 valid Entrez IDs were excluded from KEGG and Reactome analyses to ensure statistical robustness. Results were visualized using dot plots generated with the dotplot function of the clusterProfiler package, showing the top 10 enriched terms per analysis.

### Statistical Tests

Wilcoxon rank sum tests with BH multiple hypothesis correction were used for boxplot visualizations with more than 2 groups. A Student’s T-test was used for comparisons between 2 groups with assumed normalized distribution (expression data). Wilcoxon rank sum tests between 2 groups were performed when the data was not assumed to be normally distributed. The threshold for significance for all boxplot visualizations is alpha < 0.05. For correlation and linear regression analyses with more than one variable per cancer type, BH multiple hypothesis correction was used to control for false discovery rate.

### Plot Visualizations

All R-based results were visualized using ggplot^117^, ggpubr^118^ and ComplexHeatmap^119^ R packages. All biological and methods schematics were created with BioRender.com.

## ACKNOWLEDGEMENTS

This work utilized the computing resources at the NYU Grossman School of Medicine High Performance Computing (HPC) Facility and was supported by NIH P01AG051449 (D.F) and NIH/NCI U24CA210972 (D.F). T.D. and P.M. were supported by grants from NIH (R37 R37CA248631, R01 R01HG012590, and R01 R01DK135089), the Pershing Square Sohn Cancer Prize and an NFCR grant.

---

**Supplemental Figure 1.**
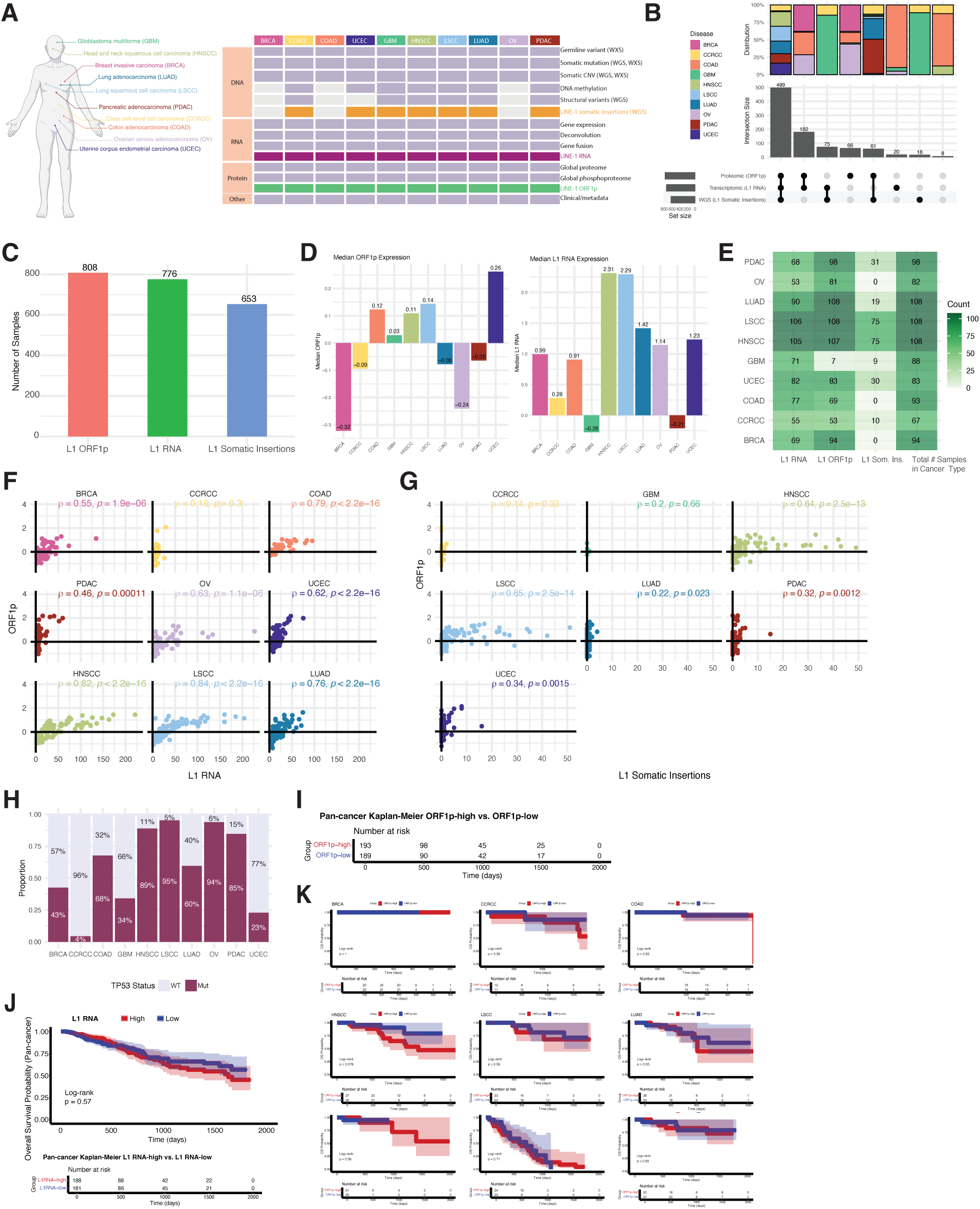
Overview of L1 across CPTAC samples. (A) Left: Schematic of CPTAC cancer tissue origin. Right: Heatmap of data type availability across CPTAC cancer types. (B) Upset plot counting number of samples containing L1 ORF1p, RNA, and insertion data across cancer types. Proportion bar plot visualizes distribution of cancer types for each set. 499 samples had quantifications in all three L1 measurements. (C) Bar plot summarizing the number of samples with each L1 measurement (n denoted above bar). (D) Bar plot summarizing the median ORF1p (left) and L1 RNA (right) expression level for each cancer type (median denoted in or above bar). (E) Heatmap summarizing the number of samples with each L1 measurement for each cancer type and the total number of samples analyzed for L1 for each cancer type. (F) Spearman correlations of L1 ORF1p (Log2 ratio) vs. L1 RNA (FPKM) per cancer type. (G) Spearman correlations of L1 ORF1p vs. L1 somatic insertion counts per cancer type. r = Spearman rho coefficient, p = p-value (H) Proportion bar plot summarizing TP53 mutant or WT status of samples per cancer type. Proportion denoted as percentage. (I) Pan-cancer sample numbers at risk for ORF1p-high and ORF1p-low groups pertaining to survival curve in Figure 1D. (J) Pan-cancer Kaplan-Meier plot comparing OS for patients stratified by L1 RNA status, log rank test. Numbers in parentheses represent the pan-cancer sample size for each group. (K) Individual Kaplan-Meier analyses for ORF1p-high vs. ORF1p-low samples per cancer type.

**Supplemental Figure 2.**
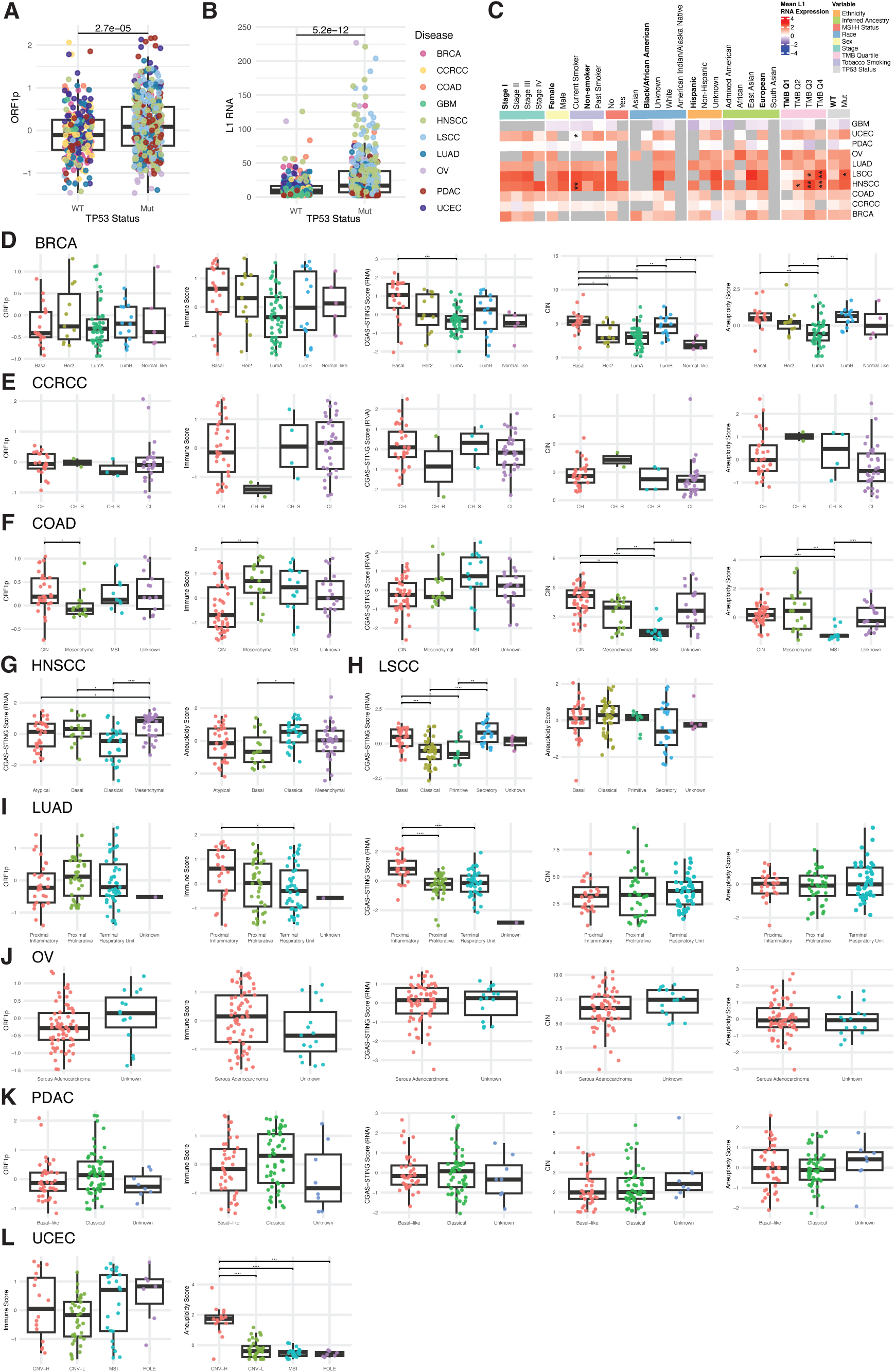
TP53, clinical variable, and ORF1p expression stratification by immune score, CIN and aneuploidy scores. (A-B) Pan-cancer ORF1p (A) and L1 RNA (B) levels by p53 status. Student’s T-test for significance. Cancer type colors for both box plots indicated on the right of (B). (C) Mean L1 RNA expression stratified by clinical variable sub-categories. Columns are sub-categories grouped by clinical variables. Rows are cancer types. Bolded sub-categories indicate the reference group used for Wilcoxon rank sum comparison of L1 RNA distribution with BH p-value correction for variables with more than 2 sub-categories for each clinical variable. Significant comparisons indicated with asterisk (*). (D-F, I-K) ORF1p expression, cytotoxic immune score (RNA-based), CGAS-STING score (RNA-based), CIN and aneuploidy score stratified by genomic subtypes per cancer type. (G-H, L) CGAS-STING scores (G-H) and cytotoxic immune score (L) and aneuploidy scores (G-H, L) for HNSCC, LSCC and UCEC, respectively. Pairwise Wilcoxon rank sum tests (BH p-value correction) were performed to test for significant comparisons. p ≤ 0.05, **p≤ 0.01, ***p ≤ 0.001, ****p ≤ 0.0001.

**Supplementary Figure 3.**
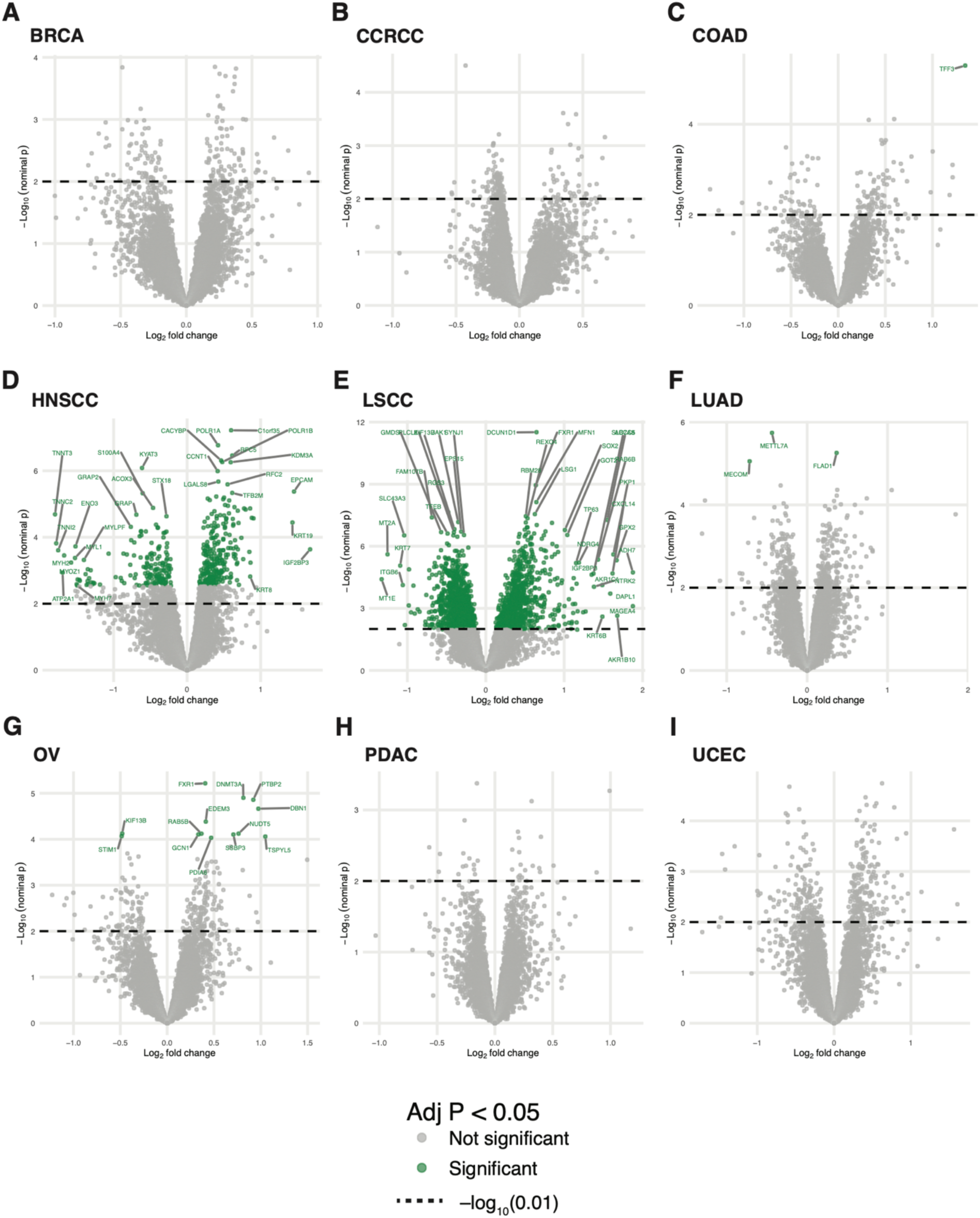
Volcano plots from DEqMS differential protein expression. (A-I) Representative volcano plots from differential proteomic expression analysis plotting the -Log10 of the nominal (raw) p-value versus the Log2 fold change (logFC) of expression. Horizontal dotted line indicates a nominal p-value of < 0.01. Significant logFCs based on the BH-adjusted p-value threshold < 0.05 are colored in green and labeled with the gene name.

**Supplementary Figure 4.**
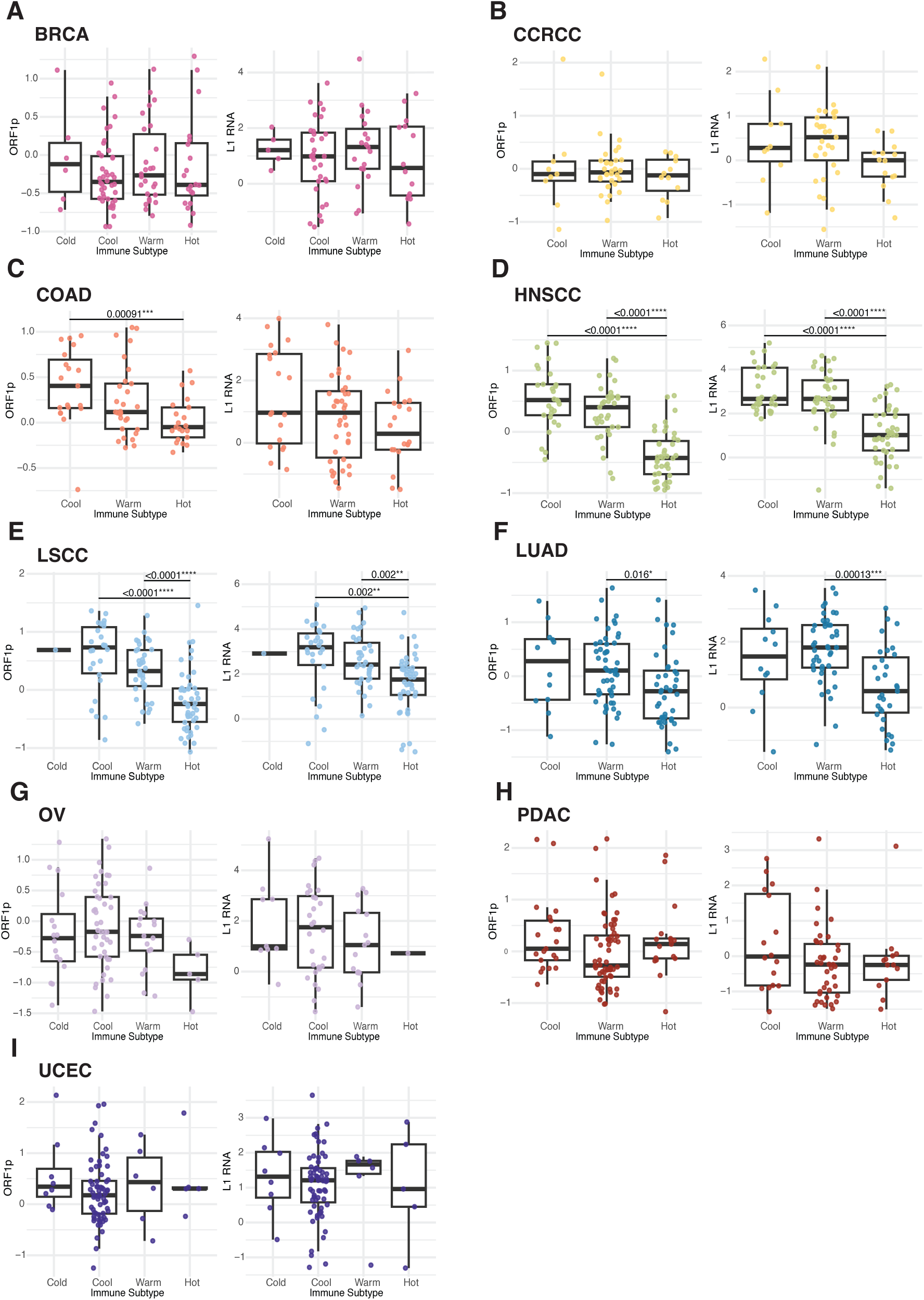
ORF1p and L1 RNA expression by CPTAC pan-cancer immune subtypes. (A-I) Box plots showing ORF1p (left) and L1 RNA (right) expression stratified by CPTAC pan-cancer immune subtypes within individual cohorts. Two-sided Wilcoxon rank sum tests (BH-adjusted P values) are performed to test for significant comparisons. p ≤ 0.05, **p≤ 0.01, ***p ≤ 0.001, ****p ≤ 0.0001.

**Supplementary Figure 5.**
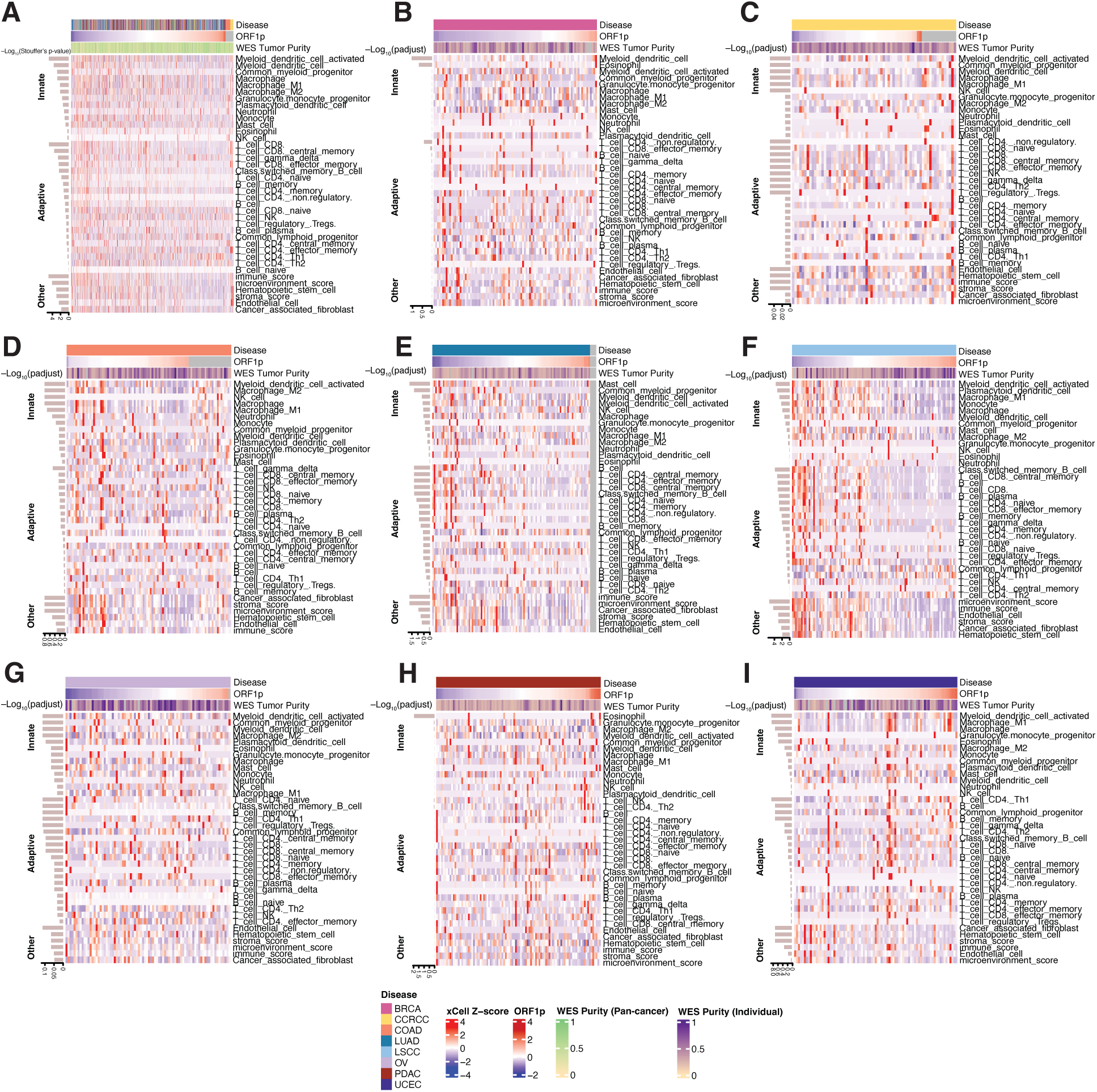
Associations of immune cell types with ORF1p. (A) Pan-cancer heatmap of xCell immune cell type deconvolution scores ordered by increasing ORF1p expression across all cancer types. Correlation results between ORF1p and xCell scores were compiled for each cancer type from previously computed linear regressions. For each immune cell type, we combined the per-cancer p-values across all remaining cancer types using Stouffer’s method, a z-score-based p-value combination approach. Rows are immune cell types grouped by branches of the immune system. Rows within each group are ordered by decreasing significance of the combined Stouffer’s P value of the ORF1p term. Whole exome purity (WES) scores denoted by yellow to green scale in legend for pan-cancer analysis. (B-I) Individual cancer type heatmaps of xCell immune cell type deconvolution scores ordered by increasing ORF1p expression. WES tumor purity covariate, colored yellow to purple in legend, also annotated. Rows are immune cell types grouped by branches of the immune system. Rows within each group are ordered by decreasing significance of the P adjusted value of the ORF1p term from the linear regression model.

**Supplemental Figure 6.**
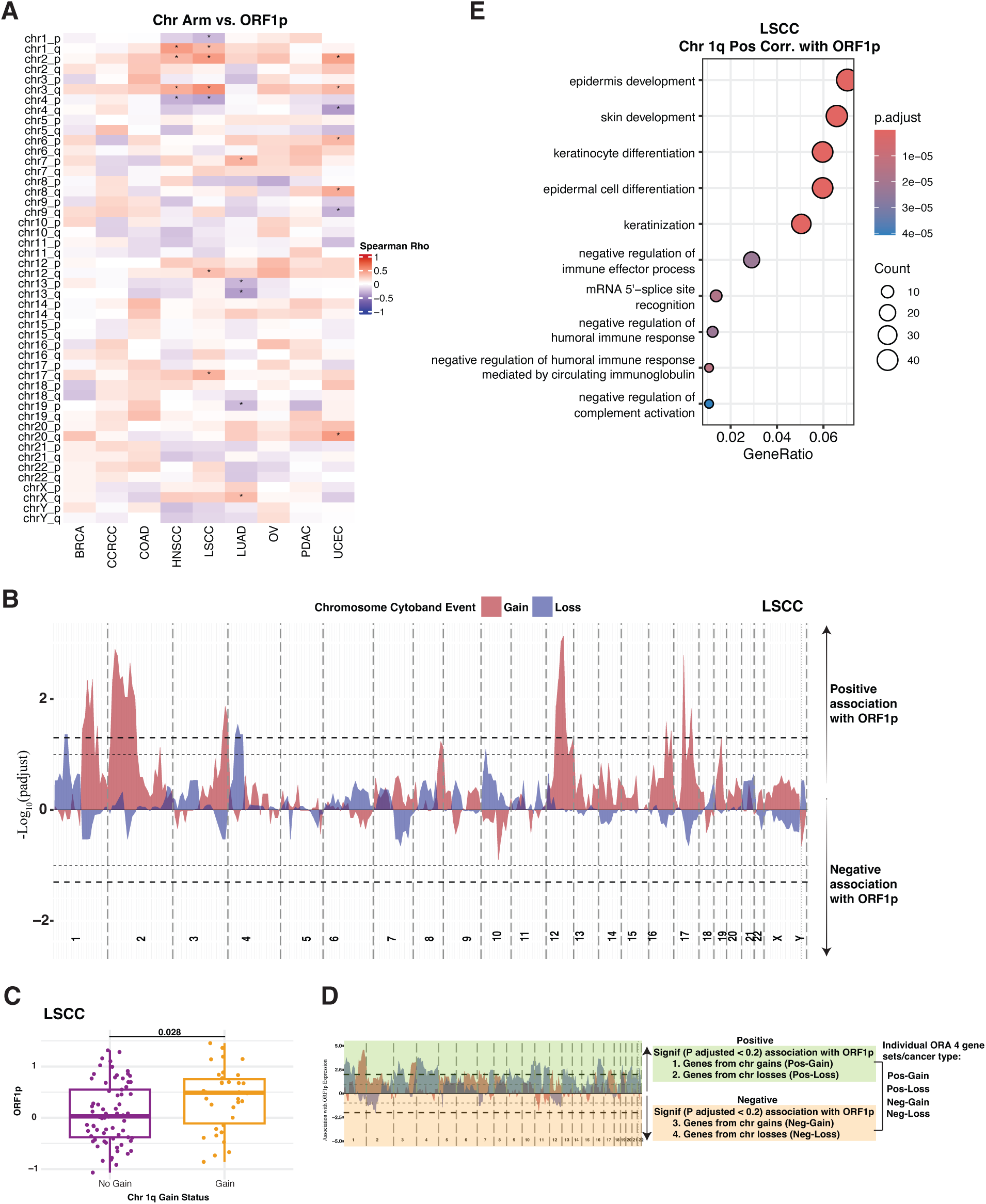
Associations of ORF1p expression with copy number alterations. (A) Heatmap of Spearman correlation coefficient (rho) between chromosome arm (row) copy number ratio and ORF1p expression per cancer type (columns). For each chromosomal arm, we computed Spearman’s rank correlation between its copy number ratio and ORF1p expression. Multiple testing correction was performed using the BH method to obtain adjusted p-values per cancer type. Significance (adjusted P value < 0.05) is annotated (*). (B) Area plot represents the association between chromosome cytoband gains (red) and losses (blue) and ORF1p expression in LSCC. Chr 1q is boxed. (C) ORF1p expression in LSCC is stratified by gain or no gain of 1q. Student’s T-test for significance. (D) Schematic outlining ORA workflow for gained (red) or lost (blue) genes significantly associated with ORF1p expression in the positive (green) or negative (orange) direction. ORA was performed for each gene set per cancer type. (E) ORA dot plot for ORF1p-predicted (adjusted P < 0.05) gained 1q cytobands in LSCC. Dot plot is showing top 10 Gene Ontology Biological Pathways (GOBP) from ORA for all genes located on ORF1p-predicted (adjusted P < 0.05) gained 1q cytobands in LSCC.

